# Modulation of GPR133 (ADGRD1) Signaling by its Intracellular Interaction Partner Extended Synaptotagmin 1 (ESYT1)

**DOI:** 10.1101/2023.02.09.527921

**Authors:** Gabriele Stephan, Hediye Erdjument-Bromage, Wenke Liu, Joshua D. Frenster, Niklas Ravn-Boess, Devin Bready, Julia Cai, David Fenyo, Thomas Neubert, Dimitris G. Placantonakis

## Abstract

GPR133 (ADGRD1) is an adhesion G protein-coupled receptor that signals through Gαs and is required for growth of glioblastoma (GBM), an aggressive brain malignancy. The regulation of GPR133 signaling is incompletely understood. Here, we use proximity biotinylation proteomics to identify ESYT1, a Ca^2+^-dependent mediator of endoplasmic reticulum-plasma membrane bridge formation, as an intracellular interactor of GPR133. ESYT1 knockdown or knockout increases GPR133 signaling, while its overexpression has the opposite effect, without altering GPR133 levels in the plasma membrane. The GPR133-ESYT1 interaction requires the Ca^2+^-sensing C2C domain of ESYT1. Thapsigargin-mediated increases in cytosolic Ca^2+^ relieve signaling-suppressive effects of ESYT1 by promoting ESYT1-GPR133 dissociation. ESYT1 knockdown or knockout in GBM impairs tumor growth *in vitro*, suggesting functions of ESYT1 beyond the interaction with GPR133. Our findings suggest a novel mechanism for modulation of GPR133 signaling by increased cytosolic Ca^2+^, which reduces the signaling-suppressive interaction between GPR133 and ESYT1 to raise cAMP levels.

## Introduction

The 33 members of adhesion family of G protein-coupled receptors (aGPCRs) have recently received attention for their roles in both physiological processes, as well as disease (Einspahr and Tilley, 2022; Hamann et al., 2015; Kaczmarek et al., 2021; Krishnan et al., 2016; Langenhan, 2020; Scholz et al., 2019). Adhesion GPCRs are characterized by an intracellular C-terminus, seven transmembrane segments and long extracellular N-termini that adhere to components of the extracellular matrix or membrane proteins. These N-termini contain receptor-specific domains that determine binding to extracellular interactors, as well as a conserved GPCR Autoproteolysis Inducing (GAIN) domain, which catalyzes autoproteolytic cleavage at the GPCR Proteolysis Site (GPS) to generate N-terminal (NTF) and C-terminal (CTF) fragments (Arac et al., 2012; Frenster et al., 2021). Immediately distal to the GPS lies the *Stachel* sequence, which has been shown by functional and structural data to act as an endogenous tethered agonist, by binding an orthosteric binding groove with the 7-transmembrane portion of aGPCRs (Barros-Alvarez et al., 2022; Liebscher et al., 2014; Ping et al., 2022; Qu et al., 2022; Xiao et al., 2022). Several lines of evidence suggest that NTF-CTF dissociation may promote *Stachel*-dependent receptor activation, although it is not absolutely necessary (Frenster *et al*., 2021).

Signaling mechanisms entrained by aGPCRs are receptor-specific and may also vary as a function of tissue and biological context. While some aGPCRs exhibit promiscuity in their coupling to G proteins, others demonstrate a dominant predilection for specific G proteins. Less is known about other intracellular regulatory mechanisms that influence trafficking to the cell membrane, signaling, and possible endocytosis, recycling or desensitization.

We recently demonstrated that GPR133 (ADGRD1), a member of the aGPCR family, is crucial for tumor growth in glioblastoma (GBM), an aggressive primary brain malignancy (Bayin et al., 2016; Frenster et al., 2017; Frenster et al., 2020; Frenster *et al*., 2021; Stephan et al., 2022). GPR133 couples to Gαs to increase intracellular cAMP (Bianchi et al., 2021; Bohnekamp and Schoneberg, 2011; Frenster *et al*., 2021; Liebscher *et al*., 2014; Stephan *et al*., 2022). Our previous work has helped elucidate mechanisms of activation of GPR133 (Frenster *et al*., 2021; Stephan *et al*., 2022). In particular, we proposed that NTF-CTF dissociation promotes receptor signaling, even though an uncleavable GPR133 mutant (H543R) is still signaling-competent, albeit at mildly reduced levels relative to wild-type GPR133 (Frenster *et al*., 2021). GPR133 manifests elevated basal levels of Gαs-mediated signaling (Bianchi *et al*., 2021; Bohnekamp and Schoneberg, 2011; Frenster *et al*., 2021; Stephan *et al*., 2022); however, the intracellular mechanisms that regulate GPR133 signaling are not understood. Here, we use proximity biotinylation proteomics to identify intracellular interactors of GPR133. We demonstrate that ESYT1 (Extended Synaptotagmin 1), an endoplasmic reticulum (ER)-associated protein that forms ER-plasma membrane (PM) tethers in Ca^2+^-dependent manner (Bian et al., 2018; Chang et al., 2013; Fernandez-Busnadiego et al., 2015; Ge et al., 2022; Giordano et al., 2013; Idevall-Hagren et al., 2015; Kang et al., 2019; Min et al., 2007; Saheki et al., 2016; Schauder et al., 2014), interacts with GPR133 to attenuate Gαs-mediated signaling in both HEK293 and patient-derived GBM cells. This protein-protein interaction does not regulate trafficking of GPR133 to the PM. The ESYT1-mediated attenuation of GPR133 signaling depends on a Ca^2+^-sensing C2 domain of ESYT1 and is relieved by increases in intracellular Ca^2+^. ESYT1 knockdown or knockout in patient-derived GBM cells impairs tumor growth *in vitro*, suggesting that ESYT1 has other functions beyond regulation of GPR133 signaling. Overall, our findings suggest that ESYT1, a novel intracellular interactor of GPR133, regulates GPR133 signaling in Ca^2+^-dependent manner. This observation links Ca^2+^ flux to GPR133 signaling via Gαs and cAMP, and establishes a paradigm for regulation of aGPCR activation by other cellular signaling mechanisms.

## Results

### Identification of ESYT1 as intracellular interactor of GPR133

To identify intracellular interactors of GPR133, we used an approach combining proximity biotinylation, affinity purification and mass spectrometry (**Fig. 1A**). We transfected HEK293T cells with GPR133 fused to a C-terminal BioID2 (Kim et al., 2016), which catalyzes biotinylation of intracellular proteins in close proximity to GPR133. In addition to the wild-type receptor (GPR133-BioID2), we included a cleavage-deficient and signaling-impaired double mutant GPR133 (H543R/T545A-BioID2) in our study, in order to identify both signaling-dependent and - independent interactors (Frenster *et al*., 2021; Ping *et al*., 2022; Qu *et al*., 2022). Mutation of H543 to R prevents cleavage of GPR133 into NTF and CTF and mildly reduces receptor signaling (Frenster *et al*., 2021). An additional mutation at T545, representing the first residue of the endogenous *Stachel* sequence, abolishes GPR133 signaling. Overexpression of both constructs in transfected HEK293T cells was confirmed by immunofluorescent staining and immunoblot (**Suppl. Fig 1A**,**B**). After treatment with biotin, immunoblot for biotin indicated self-biotinylation of the BioID2 fusions (**Suppl. Fig. 1B**). The signaling capacity of BioID2-fused GPR133 was confirmed by quantifying cAMP levels using a homogeneous time resolved fluorescence (HTRF)- based assay (**Suppl. Fig 1C**). Intracellular cAMP levels were significantly increased in cells overexpressing wild-type GPR133-BioID2, compared to the vector control or the signaling-deficient mutant H543R/T545A-BioID2.

**Figure 1:**
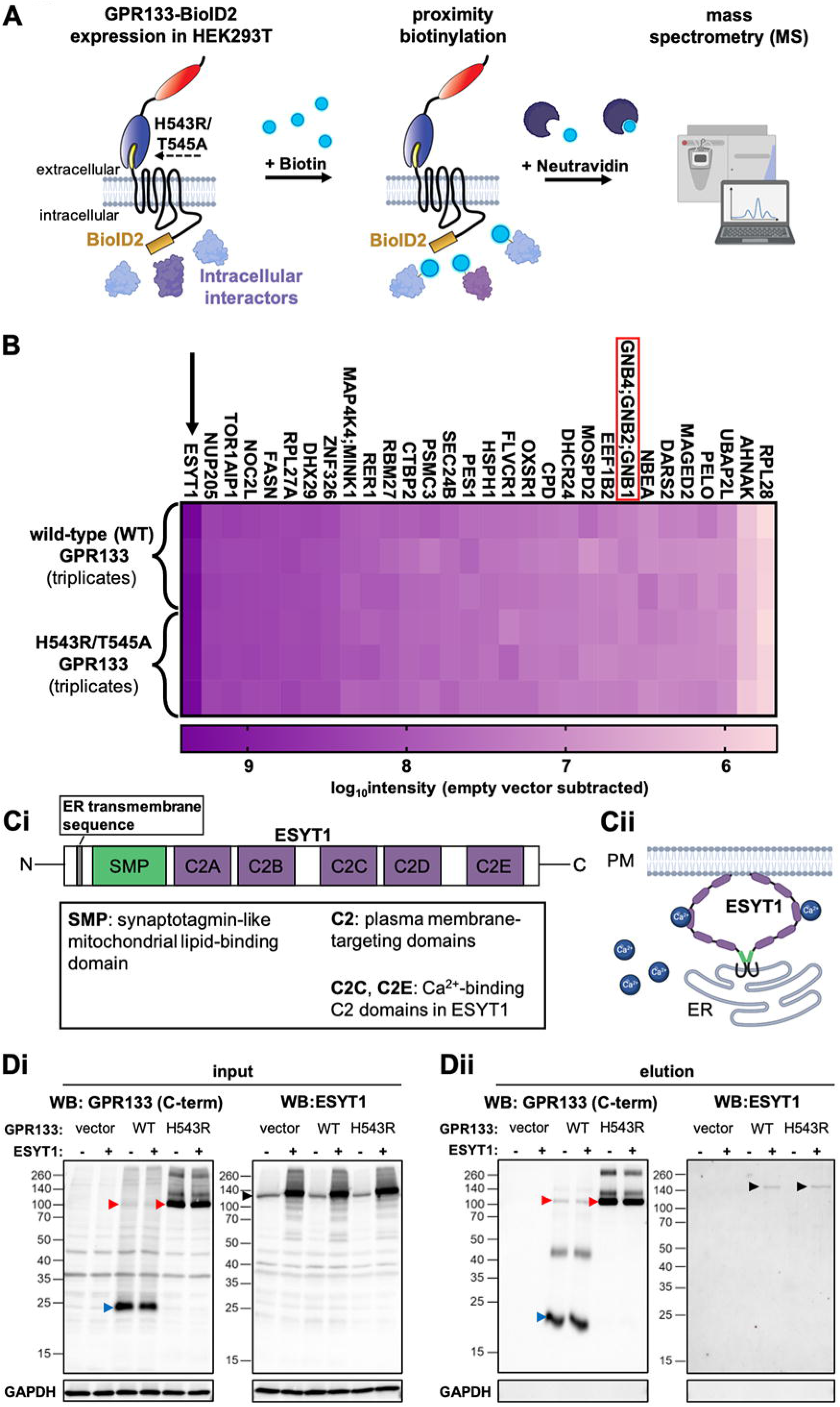
Identification of ESYT1 as a novel cytosolic interaction partner of GPR133. (**A**) Experimental design: BioID2-fusion constructs of wild-type (WT) or mutant (H543R/T545A) GPR133 were overexpressed in HEK293T cells. Following treatment with biotin, biotinylated proteins were purified using Neutravidin beads. Purified proteins were analyzed by mass spectrometry. (**B**) Top 30 biotinylated proteins with statistically equivalent detection in the two experimental conditions were ranked based on their mean MS intensity. ESYT1 (arrow) shows the highest intensity of all biotinylated proteins in close proximity to GPR133, independent on GPR133 cleavage and signaling. Gβ subunits are also identified (red box). (**C**) Structure and function of ESYT1. (**Ci**) Structural domains of ESYT1. (**Cii**) ESYT1 dimers form ER-PM tethers in response to elevations in cytosolic Ca^2+^. (**D**) Co-purification confirms binding of ESYT1 to TwinStrep-tagged GPR133, both WT and the uncleavable H543R mutant. (**Di**) Input samples: Whole cell lysates of HEK293T cells expressing wild-type GPR133 or the cleavage-deficient mutant GPR133 (H543R) with a C-terminal TwinStrep-tag following transfection with ESYT1. (**Dii**) Elution samples following Strep-Tactin purification. WB, Western blot; C-term, antibody against the cytosolic C-terminus of GPR133.

To identify intracellular GPR133 interactors, transfected HEK293T cells were treated with biotin and biotinylated proteins were purified from whole cell lysates using neutravidin beads (**Fig. 1A**). Samples were analyzed by western blot before (input) and after neutravidin purification (elution) (**Suppl. Fig 1D**). Proteins from the elution samples were further analyzed by mass spectrometry (MS) (**Fig. 1A**) (**Table 1**). Our analysis identified signaling-independent interactors that were biotinylated by both wild-type and signaling-deficient GPR133 (**Fig. 1B**) (**Table 2**), as well as signaling-dependent proteins that preferentially associated with the signaling-competent wild-type GPR133 or the signaling-deficient H543R/T545A mutant (**Suppl. Fig. 2**) (**Table 3**). In the group of shared signaling-independent interactors (**Fig. 1B**), we identified extended synaptotagmin 1 (ESYT1) as the biotinylated protein with the highest mean intensity in HEK293T cells transfected with either GPR133-BioID2 or H543R/T545A-BioID2 (arrow in **Fig. 1B**). ESYT1 belongs to a family of evolutionarily conserved proteins (ESYT1-3 in mammals) that are associated with the endoplasmic reticulum (ER) but form (ER)-to-plasma membrane (PM) tethers that mediate lipid exchange between the ER and PM without fusion of the lipid bilayers. Such lipid transfer is mediated by homo- or hetero-multimerization of members of the ESYT family through their SMP domains (Schauder *et al*., 2014) (**Fig. 1Ci**). Among the mammalian ESYT proteins, ESYT1 is unique because its ER-PM tethering function is Ca^2+^-dependent (**Fig. 1Ci**,**ii**) (Bian *et al*., 2018; Chang *et al*., 2013; Fernandez-Busnadiego *et al*., 2015; Ge *et al*., 2022; Giordano *et al*., 2013; Idevall-Hagren *et al*., 2015; Kang *et al*., 2019; Saheki *et al*., 2016). We also identified G subunit as a signaling-independent GPR133 interactor (GNB4 or GNB2 or GNB1) (red box in **Fig. 1B**), which inspired confidence in our approach. Overall, this finding raised the possibility that ESYT1 interacts with GPR133 regardless of the latter’s signaling capacity.

To validate the structural interaction between GPR133 and ESYT1, we used a co-purification approach. We generated a stable cell line of HEK293T cells expressing wild-type GPR133 (WT) or a cleavage-deficient (H543R) GPR133 mutant, both tagged with a Twin-Strep-tag at the C-terminus (Frenster *et al*., 2021). We transfected these cells with ESYT1 and prepared whole cell lysates (**Fig. 1Di**). Western blot analysis confirmed expression of GPR133 and ESYT1. As expected, using an antibody against the GPR133 cytosolic C-terminus, we detected a band corresponding to the cleaved GPR133 CTF (blue arrow, ∼25 kDa) in cells overexpressing GPR133 WT and a band representing the uncleaved full-length receptor (red arrow, ∼110 kDa) in cells overexpressing GPR133 H543R (**Fig. 1Di**) (Frenster *et al*., 2021). A minor band representing a small fraction of uncleaved receptor was also seen in cells overexpressing GPR133 WT, as expected (red arrow, **Fig. 1Di**) (Frenster *et al*., 2021). Using an antibody against ESYT1, we identified bands corresponding to the predicted molecular weight of ESYT1 (black arrows, ∼123 kDa). We were able to detect endogenous ESYT1 in empty vector-transfected cells, as well as overexpression of exogenous ESYT1 in ESYT1-transfected cells (black arrows, **Fig. 1Di**). Next, we isolated Strep-tagged WT and H543R GPR133 by affinity purification using Strep-Tactin® XT coated magnetic beads. Western blot analysis of elution samples using an antibody against the GPR133 C-terminus confirmed purification of the CTF (blue arrow, ∼25 kDa) as well as small amounts of uncleaved full-length receptor (red arrow, ∼110 kDa) from WT GPR133 - overexpressing cells and only the full-length receptor (red arrow, ∼110 kDa) in elution samples from cells overexpressing H543R GPR133 (**Fig. 1Dii**). Using an antibody against ESYT1, we detected bands corresponding to ESYT1 in elution samples of cells expressing GPR133 WT or H543R transfected with ESYT1 (black arrows, **Fig. 1Dii**). The co-purification of exogenous ESYT1 from cells expressing either WT or H543R GPR133 suggests the ESYT1-GPR133 interaction does not depend on GPR133 cleavage.

### ESYT1 downregulates GPR133 signaling

Next, we tested whether perturbation of ESYT1 levels in HEK293T cells affect GPR133 expression, trafficking to the plasma membrane or signaling. To knock down (KD) ESYT1, we used HEK293T cells stably overexpressing GPR133 or an empty vector as control, and transduced them with a lentiviral shRNA construct specific for targeting exon 8 of ESYT1 (shESYT1) or a non-specific scrambled control (shSCR) with no predicted targets in the genome. Western blot analysis confirmed reduced expression of ESYT1 following HEK293T cell transduction with shESYT1 compared to shSCR (**Fig. 2A**). Overall GPR133 expression did not change in shESYT1-transduced cells compared to shSCR-transduced cells as shown by western blot of whole cell lysates (**Fig. 2A**). Cell surface expression of GPR133 at the plasma membrane was tested with ELISA of non-permeabilized cells, using an antibody against GPR133’s N-terminus (Frenster *et al*., 2021; Stephan *et al*., 2022). GPR133 surface expression was not altered following ESYT1 KD compared to control (**Fig. 2B**). In agreement with Western blot and ELISA data, immunofluorescent staining of GPR133-overexpressing HEK293T cells showed the same overall subcellular localization pattern for GPR133 after transduction with shSCR or shESYT1 (**Fig. 2C**). While ESYT1 KD had no effect on GPR133 expression levels or plasma membrane localization, cAMP levels of GPR133-expressing cells were significantly increased following ESYT1 KD, compared to control (**Fig. 2D**).

In an alternative approach, we knocked out (KO) ESYT1 using CRISPR/Cas9. HEK293T cells were lentivirally transduced with Cas9 and gRNAs specifically targeting either ESYT1 at exon 4 or the non-essential human *Rosa26* locus. A dramatic reduction of ESYT1 protein levels in the KO condition was observed by western blot, compared to the *Rosa26* control (**Fig. 2E**). *Rosa26* KO cells and ESYT1 KO cells were then transiently transfected with GPR133 or an empty vector control and GPR133 expression was confirmed by Western blot (**Fig. 2E**, bottom panel). Notably, GPR133 expression levels in whole cell lysates were unchanged in *Rosa26* KO and ESYT1 KO cells (**Fig. 2E**). In agreement with the KD approach, GPR133 surface expression was not affected by ESYT1 KO compared to the Rosa26 control in ELISA assays (**Fig. 2F**). Using an antibody against the N-terminus of GPR133 for immunofluorescent staining, we did not observe any differences in GPR133 subcellular localization between *Rosa26* KO and ESYT1 KO cells (**Fig. 2G**). Similar to ESYT1 KD, intracellular cAMP significantly increased in GPR133-overexpressing cells following ESYT1 KO compared to the *Rosa26* condition (**Fig. 2H**). Thus, the reduction of endogenous ESYT1 levels by two different approaches increased GPR133 signaling, without changing the subcellular localization or plasma membrane levels of GPR133.

**Figure 2:**
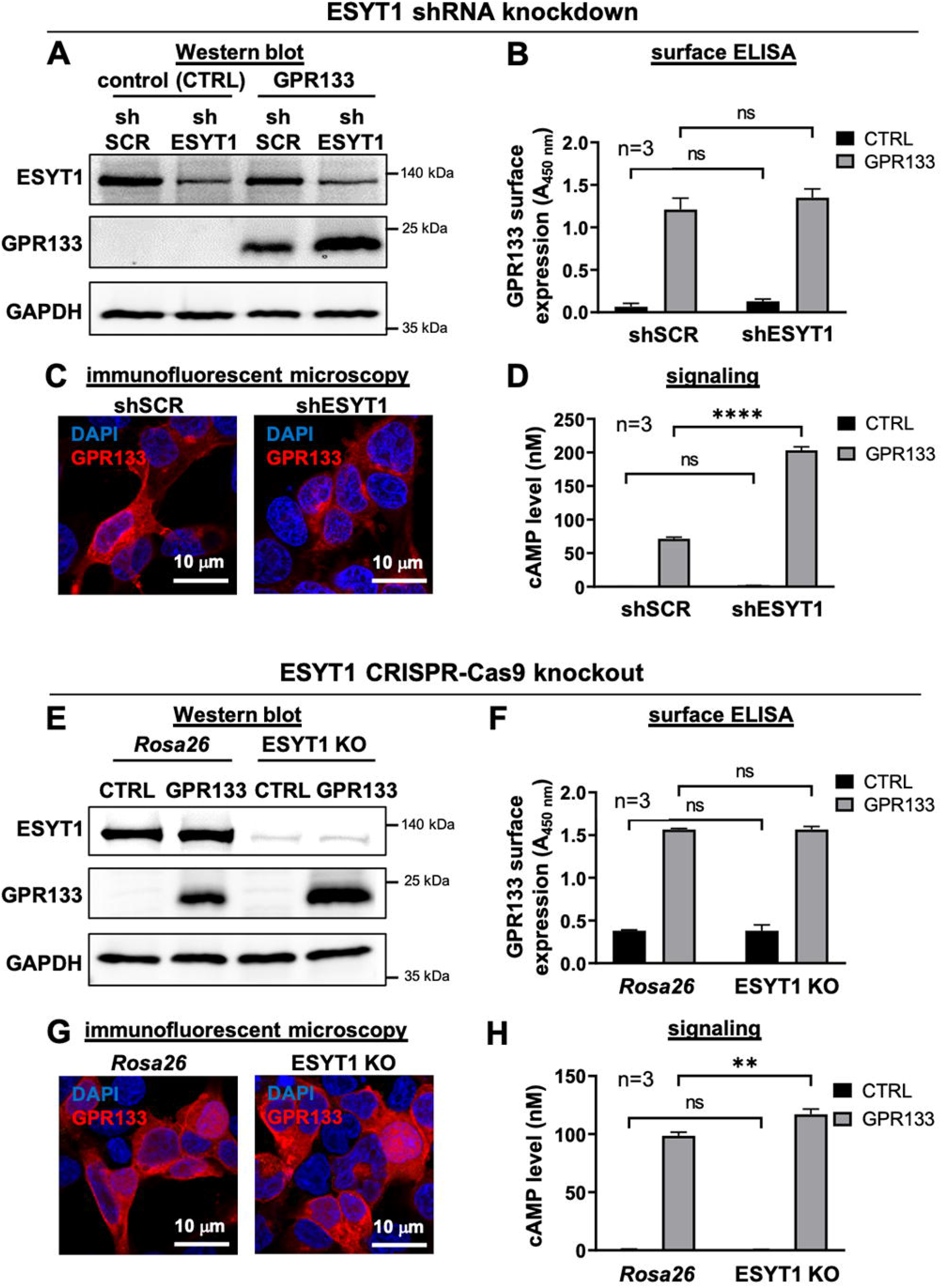
Effects of ESYT1 knockdown and knockout on GPR133 signaling. (**A-D**) ESYT1 knockdown. (**A**) Western blot confirms reduced levels of endogenous ESYT1 following its knockdown (shESYT1) compared to the control (shSCR), and stable expression of GPR133, in transduced HEK293T cells. (**B**) GPR133 surface expression is not affected by ESYT1 knockdown in ELISA assays (two-way ANOVA, p>0.05). ns, not significant; A_450 nm_, absorbance/optical density at 450 nm. Bars represent mean ± SEM of 3 experiments. (**C**) Immunofluorescent staining shows no change in the subcellular localization of GPR133 following knockdown of ESYT1 compared to the control. (**D**) Intracellular cAMP levels increase significantly in GPR133 expressing HEK293T cells after knockdown of ESYT1 compared to the control (two-way ANOVA F_(1,8)_=503.2, p<0.0001; Sidak’s *post hoc* test: GPR133 + shSCR vs. GPR133 + shESYT1, p<0.0001). Bars represent mean ± SEM of 3 experiments. (**E-H**) ESYT1 knockout. (**E**) Western blot confirms reduced levels of endogenous ESYT1 following the KO compared to the control (Rosa26). Expression of GPR133 is not affected by ESYT1 KO in transfected HEK293T cells. (**F**) GPR133 surface expression does not change upon KO of ESYT1 compared to the Rosa26 control in ELISA assays (two-way ANOVA, p>0.05). Bars represent mean ± SEM of 3 experiments. ns, not significant; A_450 nm_, absorbance/optical density at 450 nm. (**G**) Immunofluorescent staining of HEK293T cells transfected with GPR133 shows no change in GPR133 subcellular localization in ESYT1 KO cells compared to the control cells (Rosa26). (**H**) Significant increase of cAMP concentrations in GPR133 expressing HEK293T cells following KO of ESYT1 compared to the control (two-way ANOVA F_(1,8)_=10.92, p=0.0108; Sidak’s *post hoc* test: GPR133 + Rosa26 vs. GPR133 + ESYT1 KO, p=0.0027). Bars represent mean ± SEM of 3 experiments.

In a complementary approach, we tested the effects of overexpression of ESYT1 on GPR133 signaling. We generated HEK293 cells stably overexpressing an ESYT1-GFP fusion protein and transfected them with GPR133. Western blots of whole cell lysates confirmed overexpression of ESYT1 and GPR133 (**Fig. 3A**). GPR133 expression or plasma membrane localization was not affected by ESYT1 overexpression, as seen by western blot, surface ELISA and immunofluorescent staining (**Fig. 3A**-**C, Suppl. Fig. 3**). In contrast to ESYT1 KD or KO, overexpression of ESYT1 led to significant reduction in cAMP levels in GPR133-expressing cells (**Fig. 3D**).

**Figure 3:**
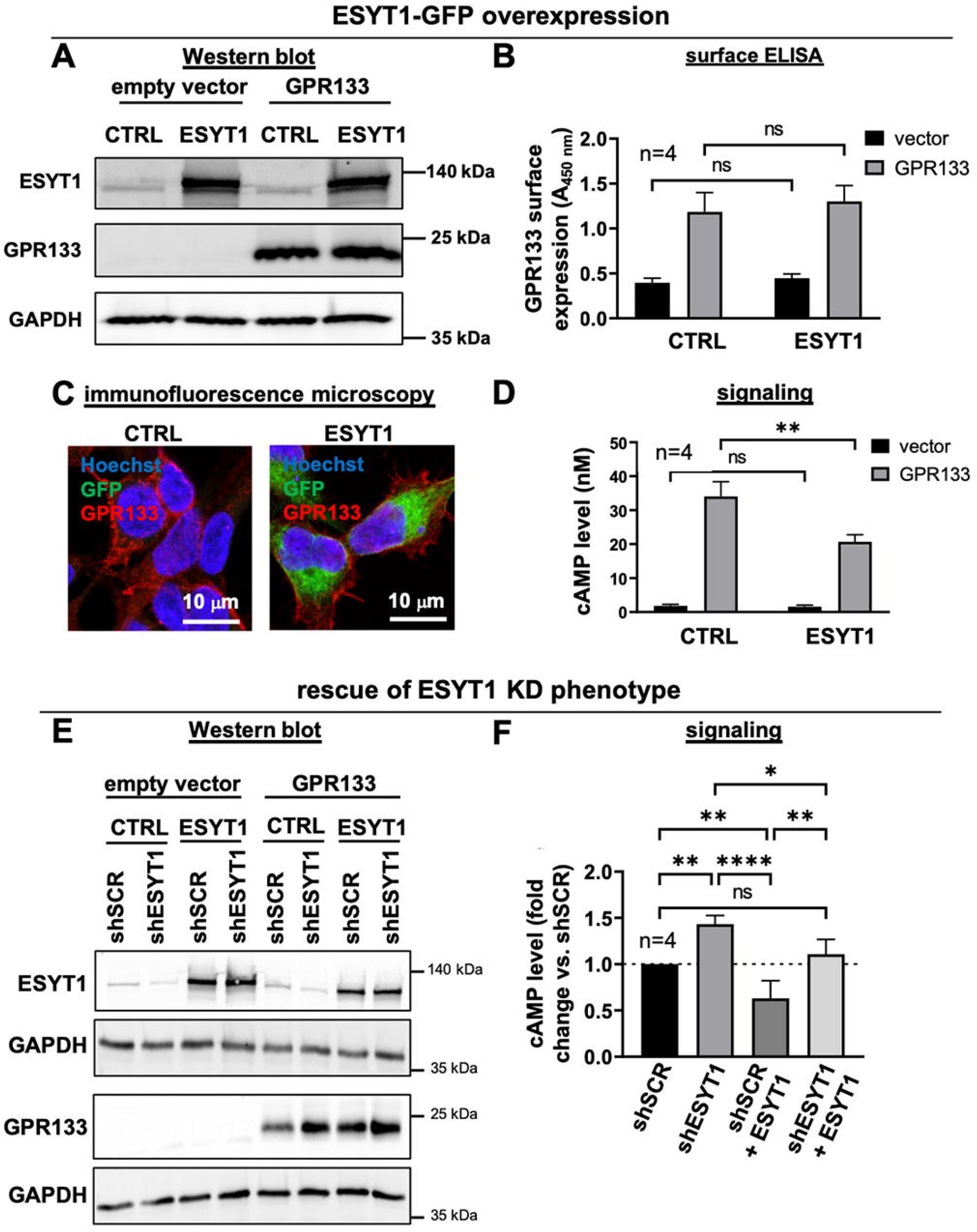
Effect of ESYT1-GFP overexpression on GPR133 signaling and expression. (**A-D**) ESYT1-GFP overexpression. (**A**) Western blot confirms increased ESYT1-GFP protein levels following transfection of GPR133 expressing cells. GPR133 expression levels are not affected in HEK293T cells. (**B**) GPR133 surface expression remains unchanged following overexpression of ESYT1-GFP (two-way ANOVA, p>0.05). Bars represent mean ± SEM of 4 experiments. ns, not significant; ns, not significant; A_450 nm_, absorbance/optical density at 450 nm. (**C**) Immunofluorescent staining of HEK293T cells expressing GPR133 and ESYT1-GFP. The cellular distribution of GPR133 immunoreactivity is unchanged. (**D**) Intracellular cAMP levels significantly decrease in GPR133-expressing HEK293T cells following overexpression of ESYT1-GFP compared to the control (two-way ANOVA F_(1,12)_=7.928, p<0.0156; Sidak’s *post hoc* test: GPR133 + CTRL vs. GPR133 + ESYT1, p=0.0041). Bars represent mean ± SEM of 4 experiments. ns, not significant. (**E-F**) ESYT1 overexpression rescues the effect of ESYT1 knockdown in GPR133-overexpressing cells. (**E**) Western Blot confirming ESYT1 knockdown and overexpression in HEK293T cells and HEK293T cells overexpressing GPR133. Expression levels of GPR133 were not affected following knockdown or overexpression of ESYT1. (**F**) Intracellular cAMP levels of GPR133 expressing HEK293T cells are normalized to shSCR. Bars represent mean ± SEM of 4 experiments. Compared to the control (shSCR), GPR133 signaling increases significantly following transduction with shESYT1 and decreases significantly following transfection with ESYT1. ESYT1 overexpression rescues the increase in cAMP levels after ESYT1 KD (one-way ANOVA F_(3,12)_=24.64, p<0.0001; Tukey’s *post hoc* test: shSCR vs. shESYT1, p=0.0030; shSCR vs. shSCR + ESYT1, p=0.0094; shESYT1 vs. shSCR + ESYT1, p<0.0001; shESYT1 vs. shESYT1 + ESYT1, p=0.0217; shSCR + ESYT1 vs. shESYT1 + ESYT1, p=0.0014). Bars represent mean ± SEM of 4 experiments. ns, not significant.

To determine whether the effects of the shRNA KD were specific to ESYT1, we tested whether overexpression of ESYT1 rescues the effect of ESYT1 KD on GPR133 signaling. We transduced HEK293T cells stably expressing GPR133 with shSCR or shESYT1 lentiviral constructs and then transfected them with exogenous ESYT1 or an empty vector control. Western blot analysis confirmed reduced ESYT1 levels in ESYT1 KD cells, but increased expression levels following transfection with ESYT1 (**Fig. 3E**). GPR133 expression levels remained unchanged throughout these perturbations (**Fig. 3E**). In HTRF signaling assays, cAMP levels increased following KD of ESYT1, compared to the shSCR condition, while they decreased following overexpression of ESYT1 (**Fig. 3F**). Importantly, ESYT1 overexpression rescued the increase in cAMP levels brought about by ESYT1 KD, suggesting specificity of the shRNA (**Fig. 3F**).

Next, we set out to determine whether the modulation of intracellular cAMP levels by ESYT1 is specific to GPR133 and not due to effects on Gαs signaling or adenylate cyclase enzymatic activity. To test whether ESYT1 may interfere with Gαs signaling, we assessed ESYT1 effects on signaling by another Gαs-coupled receptor, the 2 adrenergic receptor (ADRB2). We transduced HEK293T cells with lentiviral shSCR or shESYT1 and transfected them with a C-terminal FLAG-tagged ADRB2 or a vector control. Western blot analysis confirmed reduced ESYT1 expression in the KD condition and overexpression of FLAG-tagged ADRB2 (**Fig. 4A**). Intracellular cAMP levels of cells overexpressing ADRB2 did not change significantly following ESYT1 KD (**Fig. 4B**), suggesting the effect of ESYT1 on GPR133 signaling is specific to this receptor and not due to interference with Gαs function.

**Figure 4:**
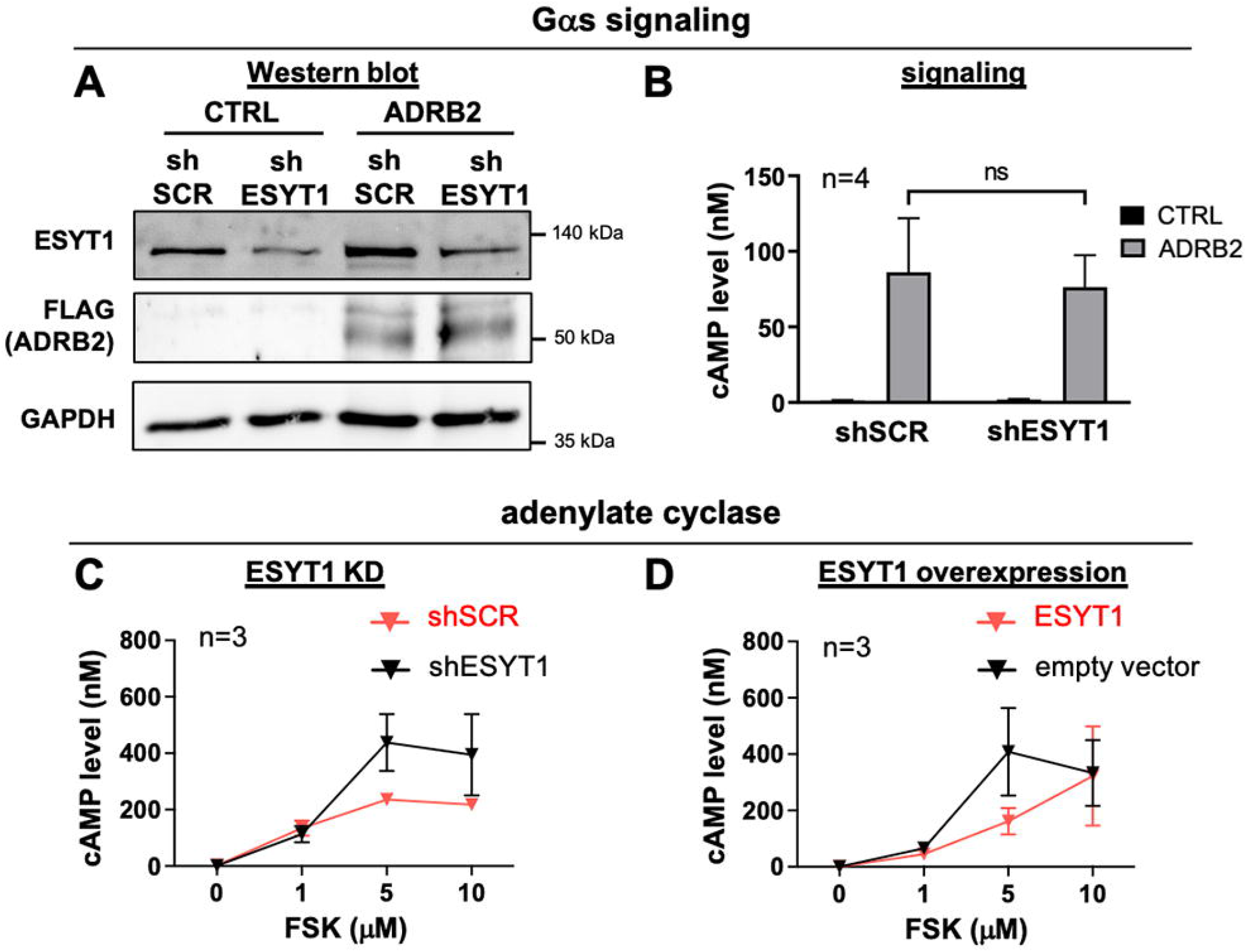
ESYT1 does not affect the Gαs-adenylate cyclase machinery. (**A-B**) Effect of ESYT1 knockdown on ADRB2 signaling. (**A**) Western blot analysis of whole cell lysates detects reduced ESYT1 levels following ESYT1 knockdown in HEK293T cells. Using a Flag-antibody, ADRB2 is detected following transfection with a Flag-tagged ADRB2 construct. (**B**) Intracellular cAMP concentrations do not change in ADRB2 expressing cells following knockdown of ESYT1 compared to the shSCR control (two-way ANOVA, p>0.05). ns, not significant. (**C-D**) Measurement of cAMP levels in HEK293T cells in response to forskolin (FSK) after either ESYT1 KD (**C**) or ESYT1 overexpression (**D**). There were no significant differences in the forskolin-induced increases in cAMP due to either perturbation in ESYT1 levels (two-way ANOVAs, p>0.05). Bars represent mean ± SEM of 2-3 experiments.

To rule out an interaction of ESYT1 with adenylate cyclase, we analyzed the effects of ESYT1 KD and ESYT1 overexpression on the cAMP response forskolin (FSK), an adenylate cyclase activator, in HEK293T cells (**Fig. 4C**,**D**). After incubation with a range of FSK concentrations (1, 5, 10 μM), intracellular cAMP levels increased indistinguishably after either ESYT1 KD (**Fig. 4C**) or ESYT1 overexpression (**Fig. 4D**), suggesting the ESYT1 levels do not influence adenylate cyclase function.

Collectively, these observations suggest that ESYT1 acts to dampen GPR133 signaling. This effect is specific to GPR133 and not due to ESYT1 interference with Gαs signaling or adenylate cyclase function.

### The ESYT1 C2C domain is required for its interaction with GPR133

ESYT1 has five C2 domains (C2A-E) (**Fig. 5A**). Among these domains, C2C and C2E are required for formation of ER-PM tethers in Ca^2+^-dependent manner (Bian *et al*., 2018; Chang *et al*., 2013; Giordano *et al*., 2013; Idevall-Hagren *et al*., 2015). The current mechanistic model posits that C2C and C2E interact at rest, but upon increases in cytosolic Ca^2+^, C2C binds Ca^2+^ to derepress C2E attachment to phospholipids in the PM, particularly PI(4,5)P_2_ [phosphatidylinositol (4,5)- biphosphate). HEK293 cells also transcribe *ESYT2* at approximately equivalent levels as *ESYT1* (Aktas et al., 2017), but ESYT2 lacks the Ca^2+^-dependence in forming ER-PM bridges and contains only three C2 domains (**Suppl. Fig. 4A**,**B**). The C2C domain of ESYT2 is considered equivalent to the C2E domain of ESYT1, in that it mediates the attachment to phospholipids in the PM (Bian *et al*., 2018; Giordano *et al*., 2013; Idevall-Hagren *et al*., 2015; Saheki *et al*., 2016). By the same token, the C2A and C2B domains are functionally conserved among all ESYT proteins and are thought to regulate the dimerization of the SMP domains, which mediate ER-PM lipid transfer (Bian *et al*., 2018). Importantly, our proximity biotinylation discovery assay suggested that the interaction of GPR133 with ESYT2 is an order of magnitude less avid than with ESYT1 (**Table 1**). Collectively, the prior knowledge and our data raise the possibility that the robust GPR133-ESYT1 interaction may be mediated by C2 domains unique to ESYT1 that confer Ca^2+^- dependence in ESYT1-mediated ER-PM bridge formation. Thus, we tested whether deletion of the C2C (ΔC2C) domain of ESYT1 modulates the interaction with GPR133 (**Fig. 5A**). We also tested whether the C2E domain, which is necessary for tethering to the PM, regulates the interaction, by deleting C2E alone (ΔC2E) or in combination with C2C (ΔC2C+E) (**Fig. 5A**).

**Figure 5:**
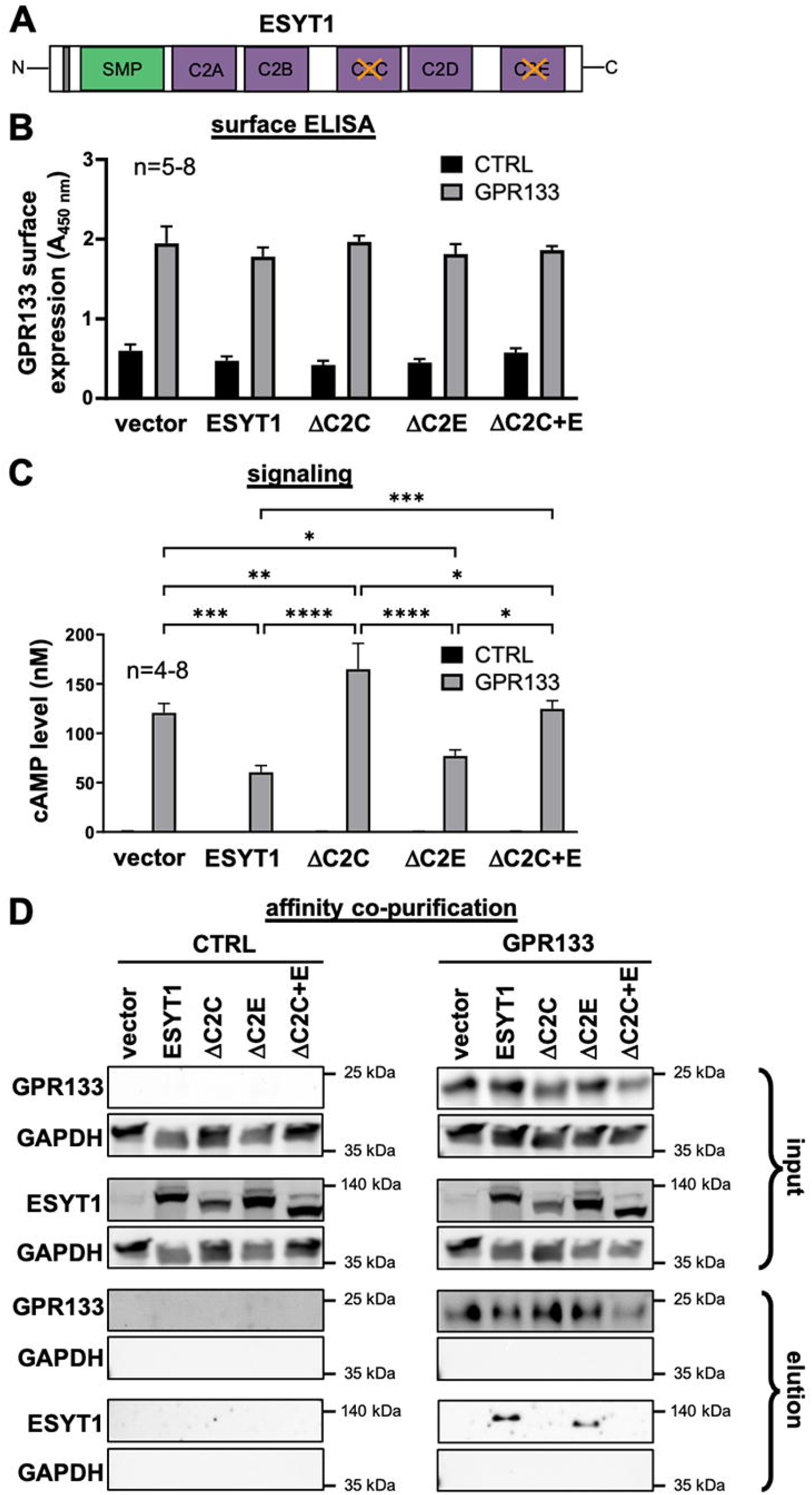
ESYT1 domains necessary for the interaction with GPR133. (**A**) Schematic showing ESYT1 deletion mutants used in this experiment. (**B**) GPR133 surface expression in ELISA assays following transfection of control HEK293T cells and HEK293T cells stably expressing GPR133 with different ESYT1 constructs. Overexpression of ESYT1, ΔC2C, ΔC2E or ΔC2C+E did not affect GPR133 surface expression compared to the vector control (two-way ANOVA, p>0.05). Bars represent mean ± SEM of 5 to 8 experiments. A_450 nm_, absorbance/optical density at 450 nm. (**C**) Intracellular cAMP levels following transfection of HEK293T cells stably expressing GPR133 with different ESYT1 wild-type or mutant constructs. Concentrations of cAMP were significantly decreased in GPR133 expressing cells after transfection with ESYT1 and ΔC2E compared to the vector control. Overexpression of ΔC2C increased cAMP levels compared to the vector control and wild-type ESYT1 in GPR133-expressing HEK293T cells (two-way ANOVA F_(4,46)_=9.471, p<0.0001; Sidak’s *post hoc* test: GPR133 + vector vs. GPR133 + ESYT1, p=0.0001; GPR133 + vector vs. GPR133 + ΔC2C, p=0.0080; GPR133 + ESYT1 vs. GPR133 + ΔC2C, p<0.0001; GPR133 + ESYT1 vs. GPR133 + ΔC2C+E, p=0.0002; GPR133 + ΔC2E vs. GPR133 + ΔC2C+E, p=0.0218). Bars represent mean ± SEM of 5 to 8 experiments. (**D**) Affinity purification analysis testing binding of different ESYT1 constructs to GPR133. Input samples represent whole cell lysates of naïve HEK293T cells and HEK293T cells stably overexpressing GPR133 transfected with ESYT1 wild-type or deletion constructs. Elution samples following Strep-Tactin purification demonstrate that ESYT1 specific bands are only detected in GPR133 expressing cells transfected with wild-type ESYT1 and ΔC2E, but not after transfection with ΔC2C or ΔC2C+E.

First, we compared the effects of all deletion mutants and full-length ESYT1 on GPR133 surface expression (**Fig. 5B**) and signaling (**Fig. 5C**). We transfected naïve HEK293T cells or HEK293T cells stably overexpressing GPR133 with wild-type ESYT1, ESYT1 ΔC2C, ESYT1 ΔC2E and ESYT1 ΔC2C+E, as well as an empty vector control. Expression of ESYT1 constructs and GPR133 was confirmed by western blots of whole cell lysates (**Suppl. Fig. 5**). GPR133 surface expression at the plasma membrane, as assessed by ELISA in non-permeabilized cells, did not change following overexpression of full-length ESYT1 or any of the deletion mutants (**Fig. 5B**). However, we found that deletion of the C2C domain, had a significant impact on GPR133 signaling (**Fig. 5C**). In contrast to the robust decrease in cAMP following transfection of wild-type ESYT1, overexpression of ΔC2C significantly increased cAMP levels in GPR133-expressing HEK293T cells compared to the empty vector control or wild-type ESYT1. The ΔC2E deletion mutant was not different from full-length ESYT1. Overexpression of the double deletion mutant ΔC2C+E had a similar positive effect on signaling as effect as the single ΔC2C deletion mutant, albeit not as pronounced. These data suggest that deletion of the C2C domain impacts the interaction between ESYT1 and GPR133. The fact that the magnitude of the effect of the ΔC2C deletion on GPR133 signaling exceeds that of the empty vector suggests that this mutant may have dominant negative effects on endogenous ESYT1, possibly mediated by heteromultimer formation between the mutant and endogenous proteins (Giordano *et al*., 2013; Schauder *et al*., 2014).

Next, we tested whether the affinity of the interaction between GPR133 and ESYT1 was affected by deletions of C2C and C2E. We performed affinity co-purification studies, using HEK293T cells stably overexpressing wild-type GPR133 with a TwinStrep-tag at the C-terminus and naïve HEK293T cells as control (CTRL). We transfected these cells with constructs for full-length ESYT1, ΔC2C, ΔC2E or ΔC2C+E, or an empty vector, and pulled down Strep-tagged GPR133 using Strep-Tactin® XT coated magnetic beads. Western blots of whole cell lysates confirmed overexpression of GPR133 and the ESYT1 constructs (**Fig. 5D**, input). The electrophoretic mobility of ESYT1 constructs reflected their predicted sizes (wild-type ESYT1 ∼123 kDa; ΔC2C ∼109 kDa; ΔC2E ∼109 kDa; ΔC2C+E ∼95 kDa). Following the Strep-Tactin-based purification, we detected GPR133 in all elution samples of cells overexpressing Strep-tagged GPR133 (**Fig. 5D**, elution, top panel). Using an antibody against ESYT1, we detected ESYT1 bands in GPR133-expressing cells transfected with either full-length ESYT1 or ΔC2E, but not ΔC2C or ΔC2C+E (**Fig. 5D**, bottom panel). Collectively, the signaling and biochemical data suggest that the ΔC2C domain of ESYT1 is essential for the interaction with GPR133.

### Modulation of GPR133 signaling by ESYT1 depends on intracellular Ca2+

ESYT1 forms ER-PM bridges in response to cytosolic Ca^2+^ flux (Bian *et al*., 2018; Chang *et al*., 2013; Fernandez-Busnadiego *et al*., 2015; Giordano *et al*., 2013; Idevall-Hagren *et al*., 2015; Kang *et al*., 2019; Saheki *et al*., 2016). C2C is the ESYT1 domain thought to be primarily responsible for the Ca^2+^ sensor properties of ESYT1 (Chang *et al*., 2013; Giordano *et al*., 2013). It was previously shown that the D724A point mutant in the C2C domain of ESYT1 renders it insensitive to Ca^2+^, thereby preventing ESYT1-mediated ER-PM bridge formation in Ca^2+^- dependent manner (Chang *et al*., 2013). To determine whether loss of Ca^2+^ sensing influences the ESYT1-GPR133 interaction, we generated the D724A mutant ESYT1 (Chang *et al*., 2013). To confirm that the D724A mutation, or deletion of C2C and C2E domains impair the Ca^2+^-induced PM localization of ESYT1, we performed confocal immunofluorescent microscopy in HEK293T cells stably overexpressing MAPPER, a fluorescent reporter for ER-PM junctions (Chang *et al*., 2013). We transfected HEK293T-MAPPER cells with Myc-tagged wild-type ESYT1, the Ca^2+^- insensitive mutant D724A, as well as the deletion mutants lacking the C2C domain, the C2E domain or both C2C and C2E domains. We compared the ESYT1 localization in these cells before and after treatment with 1 μM thapsigargin (TG). Thapsigargin increases intracellular Ca^2+^ levels by blocking its reuptake into the ER by the ER Ca^2+^ ATPase, thereby depleting Ca^2+^ stores in the ER and promoting Store-Operated Ca^2+^ Entry (SOCE) (Liou et al., 2005) (**Suppl. Fig. 6**). By staining with an antibody against GFP (detecting MAPPER, green) and Myc (detecting ESYT1, red), we detected significant increase in the overlap (orange) of wild-type, full-length ESYT1 and MAPPER following TG treatment relative to the control condition (DMSO treatment), suggesting formation of ER-PM tethers (**Fig. 6A**). However, we did not observe such overlap following TG treatment in the D724A and the deletion mutants (**Fig. 6A**). This finding suggests that translocation of ESYT1 to ER-PM bridges was prevented following the alteration of its Ca^2+^- sensing capacity.

**Figure 6:**
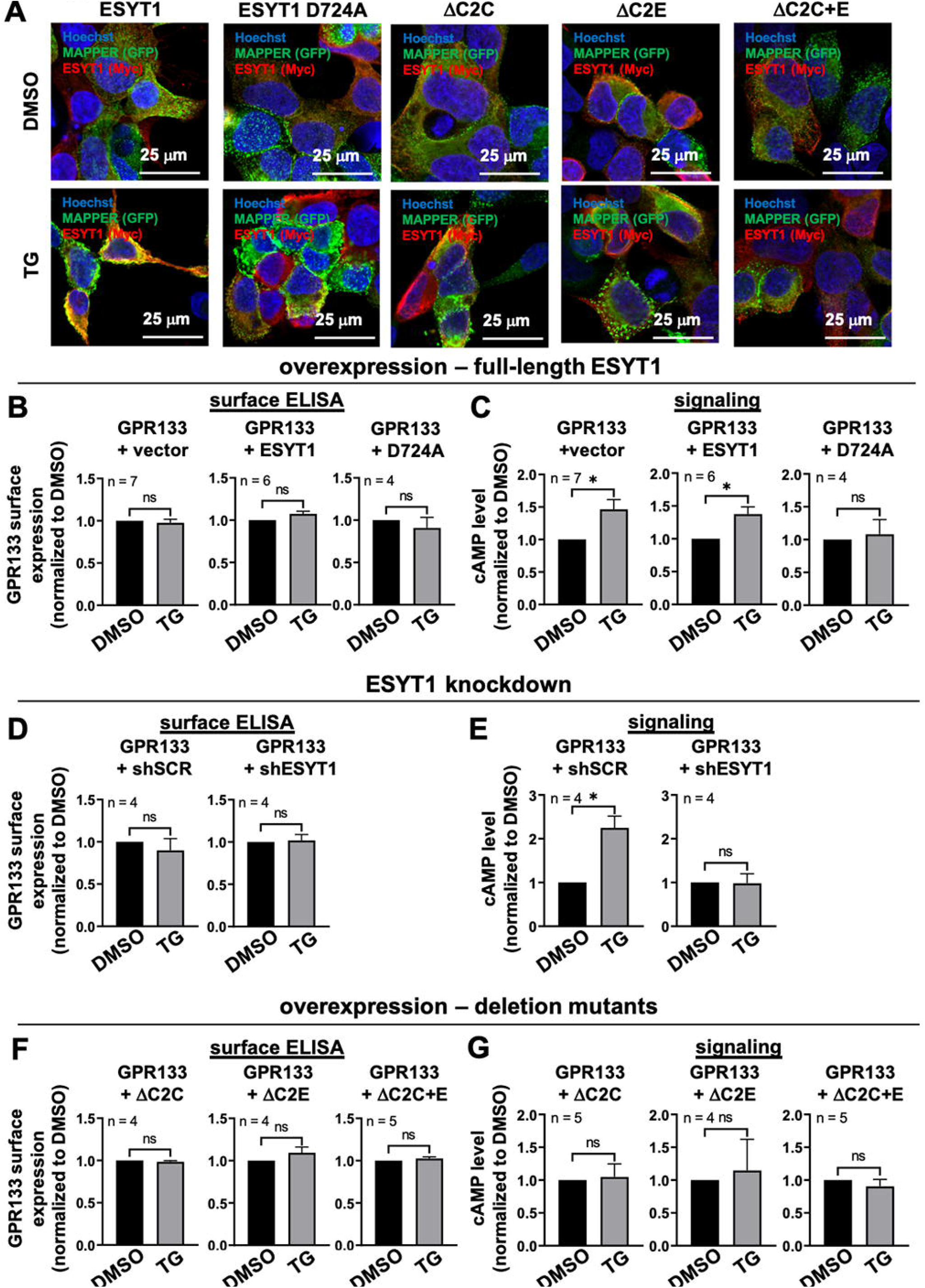
Intracellular Ca^2+^ increases impact GPR133 signaling dependent on ESYT1 expression. (**A**) Confocal images of HEK293 cells stably overexpressing MAPPER-GFP (green) transfected with Myc-tagged ESYT1 wild-type and mutant constructs (red) following treatment with DMSO or 1 μM TG to increase intracellular Ca^2+^ concentration. Yellow regions within the images represent overlap of MAPPER (green) and Myc-tagged ESYT1 (red), suggesting localization of ESYT1 at ER-PM junctions. The overlap is significantly more extensive following TG treatment of HEK293-MAPPER cells overexpressing wild-type ESYT1 rather than the mutant constructs. (**B-G**) Effect of intracellular Ca^2+^ increases on GPR133 surface expression (**B, D, F**) and cAMP levels (**C, E, G**). (**B-C**) TG Treatment of HEK293T cells stably expressing GPR133 transfected with vector, full-length ESYT1 wild-type or the mutant D724A. Bars represent mean ± SEM of 4 to 7 experiments. (**B**) TG treatment had no effect on GPR133 surface expression in GPR133 expressing HEK293T cells transfected with vector, ESYT1 or D724A compared to treatment with DMSO (paired t-test, p>0.05). (**C**) TG treatment significantly increased cAMP levels in GPR133 expressing HEK293T cells transfected with vector and ESYT1 compared to treatment with DMSO (paired t-test, GPR133 + vector: DMSO vs TG, p=0.0210; GPR133 + ESYT1: DMSO vs TG, p=0.0189). TG treatment did not affect GPR133 signaling following transfection of D724A (paired t-test, p>0.05). ns, not significant. (**D-E**) TG Treatment of HEK293T cells transduced with shSCR or shESYT1 to knockdown ESYT1. Bars represent mean ± SEM of 4 experiments. (**D**) TG treatment did not affect GPR133 surface expression compared to treatment with DMSO in GPR133 expressing HEK293T cells transduced with shSCR or shESYT (paired t-test, p>0.05). (**E**) TG treatment significantly increased cAMP concentrations compared to treatment with DMSO in HEK293T cells overexpressing GPR133 and transduced with shSCR (paired t-test, p=0.018). TG treatment had no effect on cAMP levels compared to DMSO following overexpression of GPR133 and knockdown of ESYT1 (paired t-test, p>0.05). ns, not significant. (**F-G**) TG treatment of HEK293T cells stably expressing GPR133 transfected with ESYT1 deletion mutants ΔC2C, ΔC2E or ΔC2C+E. Bars represent mean ± SEM of 4 to 5 experiments. (**F**) Treatment with TG had no effect on GPR133 surface expression in GPR133 expressing HEK293T cells transfected with ΔC2C, ΔC2E or ΔC2C+E compared to treatment with DMSO (paired t-test, p>0.05). (**G**) TG treatment did not affect cAMP concentrations compared to treatment with DMSO in GPR133 expressing HEK293T cells transfected with ΔC2C, ΔC2E or ΔC2C+E (paired t-test, p>0.05). ns, not significant.

We then tested how increases in cytosolic Ca^2+^ brought about by TG modulate GPR133 signaling in HEK293T cells stably overexpressing GPR133 and transfected with wild-type ESYT1, the Ca^2+^- insensitive mutant D724A or a vector control. D724A ESYT1 did not affect GPR133 surface expression, but decreased basal levels of GPR133 signaling, similar to the effect following transfection of wild-type ESYT1 (**Suppl. Fig. 7**). To trigger an increase in intracellular Ca^2+^, we treated cells with 1 μM TG (**Fig. 6B**,**C**). TG had no effect on GPR133 surface expression compared to control treatment with DMSO in any of the experimental groups (**Fig. 6B**). However, cAMP levels significantly increased in response to TG in cells overexpressing GPR133 and either an empty vector or wild-type ESYT1 (**Fig. 6C**). In contrast, we did not observe significant changes of cAMP levels following TG in cells transfected with the Ca^2+^-insensitive mutant D724A (**Fig. 6C**). To confirm that the Ca^2+^-dependent increase in GPR133 signaling is mediated by ESYT1, we repeated the TG treatment in HEK293T cells overexpressing GPR133 and transduced with lentiviral shSCR or shESYT1. GPR133 surface expression in ELISA assays was not affected by TG in either condition (**Fig. 6D**). While we observed significantly increased cAMP levels following the treatment of HEK293T cells expressing GPR133 and shSCR with TG compared to the DMSO control (**Fig. 6E**), the TG treatment had no effect on GPR133 signaling following ESYT1 KD (**Fig 6E**). In agreement with our previous observations, we did not detect significant changes of GPR133 surface expression (**Fig. 6F**) or TG modulation of GPR133-driven cAMP levels following overexpression of the Ca^2+^-insensitive deletion mutants ΔC2C, ΔC2E or ΔC2C+E (**Fig. 6G**).

There findings raise the possibility that ESYT1, which normally acts to dampen GPR133 signaling via an interaction mediated by its C2C domain, may dissociate from GPR133 when intracellular Ca^2+^ concentration rises, as occurs after TG treatment. To test this hypothesis, we performed a ESYT1-GPR133 proximity ligation assay (PLA) (**Fig. 7**). We transfected HEK293T cells with GPR133 alone or together with Myc-tagged ESYT1 (**Fig. 7A**). Using confocal immunofluorescent microscopy, we detected cells overexpressing GPR133 (green) or ESYT1 (red), with most of transfected cells expressing both proteins (orange arrows) (**Fig. 7A**). Western blots of whole cell lysates further confirmed overexpression of ESYT1 and GPR133 (**Fig. 7B**). We then performed the PLA assay using an anti-GPR133 (rabbit) antibody and an anti-Myc antibody (mouse) on cells transfected with either GPR133 alone or both GPR133 and ESYT1 (**Fig. 7C**). The PLA signal (red) was only detected in cells co-transfected with both GPR133 and ESYT1. Most importantly, the PLA signal weakened in cells overexpressing GPR133 and ESYT1 after treatment with TG when compared to DMSO (**Fig. 7D**). Optical sections from the imaged areas confirmed these observations (**Fig. 7E**). These findings support the hypothesis that the ESYT1-GPR133 interaction is weakened when intracellular Ca^2+^ increases.

**Figure 7:**
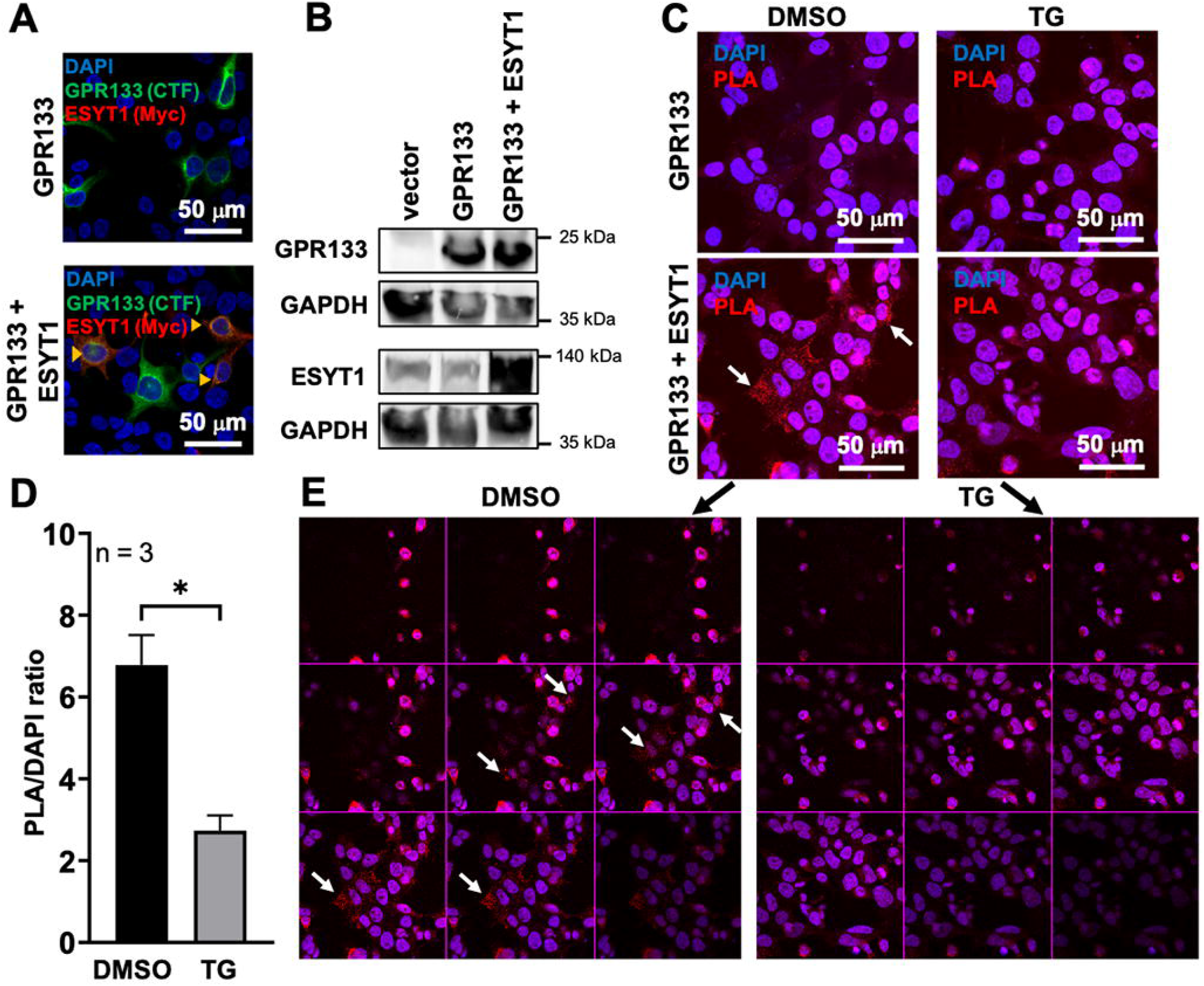
Intracellular Ca^2+^ increases disrupt binding of GPR133 and ESYT1. (**A**) Confocal images of HEK293T cells transfected with GPR133 alone (green) or co-transfected with GPR133 and Myc-tagged ESYT1 (red). In the co-transfection condition, the majority of transfected cells express both GPR133 and ESYT1 (orange arrowheads). (**B**) Western blot confirms overexpression of GPR133 and ESYT1 in transfected HEK293T cells. (**C**) Representative PLA images from in HEK293T cells transfected with GPR133 or co-transfected with GPR133 and ESYT1. The red PLA signal (arrow) is only present in cells co-transfected with GPR133 and ESYT1. The signal is weaker in cells treated with 1 μM TG compared to cells treated with DMSO. (**D**) Quantification of PLA positive signals (red dots) over DAPI positive cells overexpressing GPR133 and ESYT1. Bars represent mean ± SEM of 3 experiments. The PLA/DAPI ratio is significantly decreased in TG treated cells (paired t-test, p<0.05). (**E**) Optical sections of GPR133+ESYT1 images from the lower panel in (**C**), detecting a strong PLA signal in DMSO-treated cells (arrow), but a weaker signal in TG-treated cells.

Collectively, our data suggest that ESYT1 binds GPR133, an interaction dependent on the C2C domain of ESYT1, until cytosolic Ca^2+^ increases release ESYT1 from GPR133, thus derepressing GPR133 signaling.

### ESYT1 knockdown increases GPR133 signaling in patient-derived GBM cells

We have previously shown that GPR133 is *de novo* expressed in GBM relative to normal brain tissue and is necessary for GBM growth (Bayin *et al*., 2016; Frenster *et al*., 2017). We tested whether the effects of ESYT1 on GPR133 signaling in HEK293T cells are reproducible in patient-derived GBM cells. We transduced a patient-derived GBM culture (GBML109) with lentiviral GPR133 overexpression and shESYT1 constructs. Endogenous ESYT1 protein levels were reduced following transduction with lentiviral shESYT1 compared to shSCR, as shown by Western blot (**Fig. 8A**). Similar to HEK293T cells, cAMP levels significantly increased in GBML109 following ESYT1 KD compared to control (shSCR) (**Fig. 8B**). This finding demonstrates the impact of ESYT1 KD on GPR133 signaling in different cell types and points out its potential relevance in a disease-related biological context.

**Figure 8:**
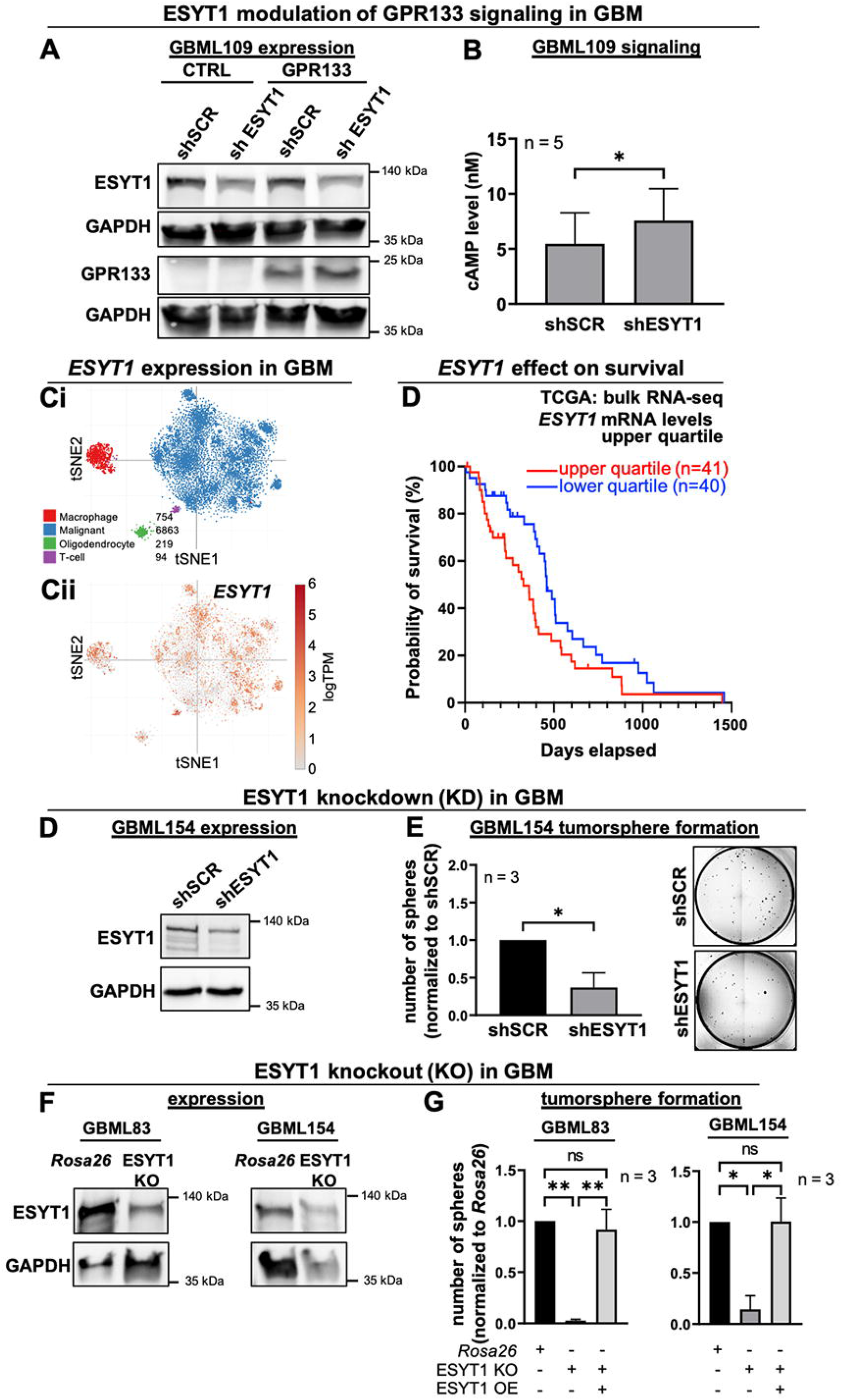
ESYT1 impacts GPR133 signaling and tumorsphere formation in patient-derived GBM cells. (**A, B**) GBML109 was transduced with lentivirus for overexpression of GPR133 and shRNA mediated knockdown of ESYT1. (**A**) Western blot analysis using specific antibodies against ESYT1 (top panel) and GPR133 (bottom panel) confirms expression of ESYT1 in GBML109 transduced with the shSCR control and knockdown of ESYT1 following transduction with shESYT1 in cells overexpressing GPR133 or an empty vector control. (**B**) Intracellular cAMP levels in GPR133-expressing GBML109 cells are significantly increased following knockdown of ESYT1 compared the control (paired t-test, p<0.05). Bars represent mean ± SEM of 5 experiments. (**C**) *ESYT1* transcript in a publically available GBM single cell RNA-seq (scRNA-seq) database (Single Single Cell Portal of the Broad Institute). (**Ci**) Identification of cellular populations in GBM specimens using tSNE plots. (**Cii**) *ESYT1* is transcribed in tumor cells, as well as macrophages, T cells and oligodendrocytes in the tumor microenvironment. (**D**) Kaplan-Meier survival curves from the TCGA GBM dataset as a function of *ESYT1* mRNA levels in bulk RNAseq of surgical specimens. Patients in the upper quartile of *ESYT1* mRNA levels experience shorter survival (median 329 days) relative to patients in the lower quartile (median 460 days) (logrank Mantel-Cox test, p=0.0413). (**E**,**F**) Effects of ESYT1 knockdown by lentiviral transduction of shRNA in GBML154. (**E**) Western blot analysis confirms KD of ESYT1 in GBML154. (**F**) Tumorsphere formation is significantly reduced in GBML154 following KD of ESYT1 compared to the control shSCR (paired t-test, p=0.0306). Bars represent mean ± SEM of 3 experiments. (**G**,**H**) Tumorsphere formation following the CRISPR/Cas9-mediated KO of ESYT1 in GBML83 and GBML154. (**G**) Reduced ESYT1 expression, detected by Western blot, following transduction with an ESYT1 specific CRISPR/Cas9 construct compared to the Rosa26 control. (**H**) Tumorsphere formation is significantly reduced in GBML83 and GBML154 following KO of ESYT1. Overexpression (OE) of ESYT1 in these cells rescues the effect (GBML83: one-way ANOVA F_(2,6)_=22.32, p=0.0017; Tukey’s *post hoc* test: *Rosa26* vs. ESYT1 KO, p=0.0023; ESYT1 KO vs. ESYT1 KO + ESYT1 OE, p=0.0036; GBML154: one-way ANOVA F_(2,6)_=10.30, p=0.0115; Tukey’s *post hoc* test: *Rosa26* vs. ESYT1 KO, p=0.0183; ESYT1 KO vs. ESYT1 KO + ESYT1 OE, p=0.0179). Bars represent mean ± SEM of 3 experiments. ns, not significant.

### Effects of ESYT1 knockdown on tumor growth *in vitro*

The role of ESYT1 in GBM has not been studied so far. We hypothesized that, If ESYT1’s principal action in GBM is to dampen GPR133 signaling, ESYT1 KD or KO should produce an increase in tumorsphere formation. However, we also anticipated that ESYT1 is likely to have additional functions in GBM, which could influence the outcome of the tumorsphere assay beyond its action on GPR133 signaling.

To determine the pattern of *ESYT1* expression in GBM, we used single cell RNA-Seq (scRNA-seq) data of adult and pediatric GBM from the Single Cell Portal of the Broad Institute (**Fig. 8Ci**). Indeed, *ESYT1* is transcribed in a significant portion of GBM cells, as well as other cell types in the tumor microenvironment (**Fig. 8Cii**). When we analyzed patient survival data as a function of *ESYT1* mRNA levels on bulk RNA-seq from surgical specimens in the GBM dataset of the TCGA (The Cancer Genome Atlas), we found that elevated *ESYT1* expression levels inversely correlate with survival (**Fig. 8D**). This raised the possibility that increased *ESYT1* expression may correlate with more aggressive tumor behavior.

To gain insight into the function of ESYT1 in tumor biology, we tested whether KD or KO of ESYT1 affects clonogenic tumorsphere formation *in vitro* (Bayin *et al*., 2016) in patient-derived GBM cultures. Such assays are a measure of the potential of single tumor cells to initiate growth and are considered a metric of the frequency of GBM stem cells *in vitro*. First, we knocked down ESYT1 in the patient-derived GBM culture GBML154 (**Fig. 8E**,**F**), using the same approach as in HEK293T cells. Western blots of whole cell lysates confirmed expression of ESYT1 and its KD (**Fig. 8E)**. We then seeded single cell suspensions of GBML154 cells transduced with shSCR or shESYT1 and counted tumorspheres that formed after two weeks (**Fig. 8F**). The number of tumorspheres was significantly reduced in GBML154 after ESYT1 KD compared to control (**Fig. 8F**). In a parallel experiment, we designed CRISPR/Cas9-mediated KO of ESYT1 in two different patient-derived GBM cultures (GBML83 and GBML154), using the same approach as in HEK293T cells. Reduction in ESYT1 protein levels was demonstrated with Western blot (**Fig. 8G**). In agreement with our findings following KD of ESYT1 in GBM cells, ESYT1 KO reduced tumorsphere formation in both GBM cultures, compared to a *Rosa26*-targeting gRNA control (**Fig. 8H**). To show the specificity of the ESYT1 KO approach, we transduced the ESYT1 KO GBM cells with lentiviral ESYT1 overexpression constructs. Indeed, the impairment in tumorsphere formation in ESYT1 KO cells was rescued following overexpression of exogenous ESYT1 (**Fig. 8H**). This finding demonstrated the specificity of the phenotype induced by the KO of ESYT1. Therefore, our experiments with GBM cells suggests that ESYT1 has multiple biological functions that extend beyond its interaction with GPR133.

## Discussion

GPR133 is an adhesion GPCR that signals through Gαs to raise cAMP, with an essential role in GBM growth (Bayin *et al*., 2016; Bianchi *et al*., 2021; Bohnekamp and Schoneberg, 2011; Frenster *et al*., 2021; Liebscher *et al*., 2014; Stephan *et al*., 2022). Previous work by us and others has demonstrated that autoproteolytic cleavage at the GPS and NTF-CTF dissociation promote receptor signaling, but are not absolutely necessary for receptor activation (Bohnekamp and Schoneberg, 2011; Frenster *et al*., 2021; Stephan *et al*., 2022). Furthermore, although ligand binding to the extracellular portion of GPR133 increases its signaling output, the receptor maintains high basal levels of activity even in the absence of ligands (Bianchi *et al*., 2021; Bohnekamp and Schoneberg, 2011; Frenster *et al*., 2021; Stephan *et al*., 2022). These facts raise the question of how cells may regulate the intrinsic signaling capacity of GPR133. Here, we have uncovered a novel interaction between GPR133 and ESYT1, an ER-anchored protein that forms ER-PM tethers via C2 domains in response to increases in cytosolic Ca^2+^ (Bian *et al*., 2018; Chang *et al*., 2013; Fernandez-Busnadiego *et al*., 2015; Ge *et al*., 2022; Giordano *et al*., 2013; Idevall-Hagren *et al*., 2015; Min *et al*., 2007; Saheki *et al*., 2016; Schauder *et al*., 2014). ESYT1 belongs to a family of three extended synaptotagmins (ESYT1-3), which all carry out the same function of tethering the ER to the PM, and by doing so, possibly mediating the exchange of phospholipids between the ER and PM lipid bilayers. However, ESYT1’s ability to form ER-PM tethers depends significantly more on Ca^2+^ than ESYT2 and 3 (Giordano *et al*., 2013).

*ESYT1* and *ESYT2* are both transcribed equivalently in HEK293T cells, in which we performed the proximity biotinylation discovery assay. However, our proteomic analysis indicated a significantly more avid interaction between GPR133 and ESYT1 relative to ESYT2. This raises the possibility that the Ca^2+^ dependence of ESYT1 may be critical in the regulation of GPR133 signaling. Indeed, the GPR133-ESYT1 interaction depends on one of the C2 domains of ESYT1, C2C, which was previously shown to be necessary for the Ca^2+^-dependent ER-PM tethering function of ESYT1 (Fernandez-Busnadiego *et al*., 2015; Ge *et al*., 2022; Giordano *et al*., 2013; Idevall-Hagren *et al*., 2015). At baseline Ca^2+^ concentrations, ESYT1 acts to suppress signaling by GPR133. However, increases in cytosolic Ca^2+^ lead to dissociation of ESYT1 from GPR133 and derepression of GPR133 signaling. This action of Ca^2+^ on GPR133 signaling is specifically mediated by ESYT1, because it is abolished when ESYT1 is knocked down or is rendered Ca^2+^- insensitive by deletion of the Ca^2+^-sensing C2 domains or the D724A point mutation in its C2C domain.

ESYT1 was previously shown to mediate PM trafficking of certain membrane proteins, such as ANO1 (Lerias et al., 2018). However, the signaling-suppressive actions of ESYT1 on GPR133 do not alter the PM localization of the latter, regardless of cytosolic Ca^2+^ levels. Several different explanations can be considered for this observation. First, it is possible that much of the GPR133 basal signaling output occurs while the receptor is being trafficked through the secretory pathway to the PM, but does not require PM localization. Second, ESYT1 may interact with PM-localized GPR133 as one component of a dynamic equilibrium that also includes the ESYT1-PM interactions, with elevations in cytosolic Ca^2+^ shifting the equilibrium to favor ESYT1-GPR133 dissociation and ESYT1-PM tethering. Finally, our ability to detect subtle changes in PM levels of GPR133 as a result of perturbation of ESYT1 levels may be technically limited.

The mechanism underlying the suppression of GPR133 signaling by ESYT1 remains unclear. GPR133 manifests elevated basal levels of Gαs-cAMP signaling, even in heterologous expression systems, such HEK293 cells, and in the absence of known ligands. This raises the possibility of baseline avid interactions with the G protein machinery. Our working hypothesis is that ESYT1 competes with G proteins for GPR133 binding. In this model, cytosolic Ca^2+^ flux derepresses signaling by reducing ESYT1 affinity for GPR133 and allowing G protein interactions with the receptor. Relevant to this hypothesis is the finding that Gβ subunits were among the top GPR133 interactors in our proximity biotinylation discovery assay. Importantly, the association of both ESYT1 and Gβ subunits with GPR133 occur independently of the signaling capacity of the receptor. In other words, both ESYT1 and Gβ subunits interact with wild-type GPR133 as avidly as they do with a mutant that is both uncleavable (H543R) and signaling-dead due to mutation of the first residue of the *Stachel* sequence (T545A). This finding suggests that the structural interactions of GPR133 with ESYT1 and Gβ do not dependent on receptor cleavage, or the potency of the endogenous orthosteric agonist *Stachel* sequence and the signaling capacity of the receptor. Future experiments will be needed to determine whether ESYT1 indeed competes with Gβ for GPR133 binding.

The GPR133-ESYT1 interaction is dynamically regulated by Ca^2+^. Increases in cytosolic Ca^2+^ lead to dissociation of ESYT1 from GPR133 on proximity ligation assays (PLA) and derepression of GPR133 signaling. This modulation represents an example of cross-talk between two dominant signaling pathways in cells: Ca^2+^ and cAMP. This mechanism may be particularly relevant in GBM, where several groups have demonstrated robust cytosolic Ca^2+^ waves in tumor cells, which promote tumor growth (Hausmann et al., 2023; Venkataramani et al., 2019; Venkataramani et al., 2022; Venkatesh et al., 2019). We postulate that such waves boost GPR133 signaling in GBM cells by causing dissociation of ESYT1 from GPR133, a mechanism that may mediate some of the tumor-promoting effects of Ca^2+^ waves and will require further testing. Interestingly, knockdown or knockout of ESYT1 in GBM reduces tumor growth *in vitro*, suggesting that ESYT1 has multiple functions in these tumors that extend beyond its interaction with GPR133.

Of particular interest is the possibility that the ESYT1-GPR133 interaction may exert bidirectional effects on the function of both proteins. Our work has uncovered effects of this interaction of GPR133 signaling, but potential regulation of ESYT1 biology by GPR133 will require future investigation. ESYT1 is thought to mediate lipid transfer between the ER and PM in Ca^2+^- dependent manner (Bian *et al*., 2018; Chang *et al*., 2013; Ge *et al*., 2022; Giordano *et al*., 2013; Idevall-Hagren *et al*., 2015; Saheki *et al*., 2016; Schauder *et al*., 2014). This action may not only be relevant to lipid metabolism, but may also help regulate signaling cascades. As an example, activation of phospholipase C (PLC) by cell surface receptors leads to hydrolysis of PI(4,5)P_2_, the PM phospholipid that the C2E domain of ESYT1 has high affinity for, to generate the second messengers inositol triphosphate (IP_3_) and diacylglycerol (DAG). The former gates IP_3_ receptors on the ER to release Ca^2+^ from ER stores into the cytosol, while the latter activates protein kinase C (PKC). Among the proposed functions of ESYT1 is the recycling of DAG from the PM to the ER, thereby potentially attenuating PKC signaling and modulating other DAG-dependent cellular processes (Xie et al., 2019). Furthermore, ESYT1 has been implicated in the regulation of SOCE (Kang *et al*., 2019). It is, therefore, plausible that GPR133 may play a regulatory role in these processes via its interaction with ESYT1.

Collectively, our findings link cytosolic Ca^2+^ flux to regulation of Gαs-cAMP signaling by GPR133 via a mechanism mediated by the GPR133-ESYT1 interaction. This is the first demonstration of cross-talk between GPR133-Gas-cAMP signaling and Ca^2+^ flux, a powerful modulator of several signaling cascades and cellular processes. We postulate that the GPR133-ESYT1 interaction will serve as paradigm for exploring similar mechanisms that modulate signaling by other aGPCRs. We theorize that this interaction may be biologically relevant to the pathogenesis of glioblastoma, a brain malignancy in which both cytosolic Ca^2+^ waves and GPR133 are essential to tumor growth. Finally, our proximity biotinylation proteomic database serves as a resource for future investigation of both “structural”, signaling-agnostic, and signaling-dependent intracellular interactors of GPR133.

## Acknowledgements

Single cell RNAseq data in GBM were obtained from the Single Cell Portal of the Broad Institute (https://singlecell.broadinstitute.org/single_cell/study/SCP393/single-cell-rna-seq-of-adult-and-pediatric-glioblastoma). Bulk RNAseq and patient survival data from the TCGA were obtained through the Xena browser (https://xenabrowser.net/). We thank the Microscopy and Flow Cytometry core facilities at NYU Grossman School of Medicine.

## Author contributions

Conceived and designed the experiments: GS, DGP. Performed the experiments: GS, HEB, WL, JDF, NR-B, DB, JC. Analyzed the data: GS, HEB, WL, DF, TN, DGP. Wrote and edited the manuscript: GS, DGP.

## Funding

This study was supported by NIH/NINDS R01 NS102665, NIH/NINDS R01 NS124920, NIH/NINDS R21 NS126806, and NYSTEM (NY State Stem Cell Science) IIRP C32595GG to DGP. DGP was also supported by NIH/NIBIB R01 EB028774, NIH/NCI R21CA263402, NIH/NCATS 2UL1TR001445, NYU Grossman School of Medicine, NYU Perlmutter Cancer Center, and DFG (German Research Foundation) FOR2149. GS was supported by a DFG postdoctoral fellowship (STE 2843/1-1). Core facilities were supported in part by Cancer Center Support Grant P30CA016087 to the Perlmutter Cancer Center of NYU Grossman School of Medicine.

## Disclosures and Competing Interests

DGP and NYU Grossman School of Medicine own an EU and Hong Kong patent titled “Method for treating high-grade gliomas” on the use of GPR133 as a treatment target in glioma. DGP has received consultant fees from Tocagen, Synaptive Medical, Monteris, Robeaute and Advantis.

## STAR Methods

### Generation of GPR133 and ESYT1 constructs

All constructs used for expression of GPR133, ESYT1 and ADRB2 or knockdown/knockout of ESYT1 are listed in **Table 4** (key resources). Primers for mutagenesis and Gibson cloning are listed in **Table 5**. Fusion constructs of wild-type GPR133-BioID2 and mutant GPR133 (H543R/T545A)-BioID2 were created using the MCS-13X Linker-BioID2-HA (addgene #80899) (Kim *et al*., 2016) plasmid and subsequently subcloned into the lentiviral backbone pLVX-EF1a-mCherry-N1 (Takara, Cat# 631986). Twin-Strep-tagged GPR133 constructs were available from previous studies (Frenster *et al*., 2021). ESYT1, ADRB2 and MAPPER cDNA plasmids were obtained from Addgene (Myc-ESYT1 #66833, EFGP-ESYT1 #66830, Flag-ADBR2 #14697, GFP-MAPPER #117721). The D724A point mutant from ESYT1 was generated by site-directed mutagenesis using the Q5® Site-Directed Mutagenesis Kit (NEB, Cat# E0554S), following the manufacturer’s protocol. ESYT1 deletion mutants Δ1C2C, Δ1C2E and Δ1C2C+E were created from the full-length ESYT1 sequence using a two-fragment Gibson reaction.

### Cell culture

Human embryonic kidney 293 T cells (HEK293T, Takara, Cat# 632180) were cultured in Dulbecco’s modified Eagle’s medium (DMEM, Gibco, Cat# 11965-118) with sodium pyruvate (Gibco, Cat# 11360070) and 10% fetal bovine serum (FBS; Peak Serum, Cat# PS-FB2) at 37 °C and 5% CO_2_. Patient-derived GBM cultures were established as previously described (Frenster and Placantonakis, 2018). GBM cells were cultured in Neurobasal medium (Gibco, Cat# 21103049) supplemented with N2 (Gibco, Cat# 17-502-049), B27 (Gibco, Cat# 12587010), non-essential amino acids (Gibco, Cat# 11140050) and GlutaMax (Gibco, Cat# 35050061), as well as 20 ng/mL recombinant basic Fibroblast Growth Factor (bFGF; R&D, Cat# 233-FB-01M) and 20 ng/mL Epidermal Growth Factor (EGF; R&D, Cat# 236-EG-01M) at 37°C, 5% CO_2_ and 4% O_2_. GBM cells were grown in spheroid suspensions or as attached cultures on cell culture dishes, pretreated with poly-L-ornithine (Sigma, Cat# P4957) and laminin (Thermo Fisher, Cat# 23017015). HEK293T cells were passaged using Trypsin (Thermo Fisher, Cat# 25300054) and GBM cells were dissociated and passaged with Accutase (Innovative Cell Technologies, Cat# AT104).

### Transfection and lentiviral infection

HEK293T cells were transfected with plasmid DNA using Lipofectamine 2000 (Invitrogen, Cat# 11668-019), following the manufacturer’s protocol. Selection of HEK293T cells stably transfected with ESYT1-GFP was performed with G418 (ThermoFisher, Cat# 10131035). HEK293T or GBM cells were transduced using lentivirus as described previously (Frenster et al., 2018). In short, lentivirus was produced by co-transfecting HEK293T cells with expression plasmids of interest and packaging plasmids psPax2 and pMD2.G. Lentivirus was collected from the cell culture supernatant 24h, 48h, and 72h after transfection and concentrated using the Lenti-X concentrator (Contech Takara, Cat# 631231). For lentiviral transduction, HEK293T or GBM media was supplemented with 4 μg/ml protamine sulfate and cells were treated with viral particles at a multiplicity of infection (MOI) of three. Stable cell lines were selected with 5 μg/mL puromycin and by isolating mCherry- or GFP-positive cells by fluorescence-activated cell sorting (FACS) with the SH800Z sorter (Sony Biotechnology).

### Western blot analysis of whole cell lysates

Cells were washed with PBS and lysed in RIPA buffer (Thermo, Cat#89900) containing Halt protease inhibitor cocktail (Thermo, Cat# 78429) and 1% n-dodecyl β-D-maltoside (DDM) (Thermo, Cat# BN2005). Lysates were sonicated in a water-bath Bioruptor (Diagenode, Cat# UCD-300) and precleared at 15,000 x g for 10 min at 4 °C. Protein concentrations were measured using the DC protein assay kit II (BioRad, Cat# 5000112). Laemmli buffer (BioRad, Cat# 1610747) containing DTT (BioRad, Cat# 1610610) was then added and samples were incubated at 37 °C for 30 min. Twenty μg of protein samples were separated by SDS-PAGE and transferred to nitrocellulose membranes (BioRad, Cat# 1620112). Membranes were blocked in 2% bovine serum albumin (BSA) in TBS-Tween for 1 hour at room temperature (RT), incubated with primary antibodies (**Table 4**) at 4 °C overnight, washed with TBS-Tween and incubated with Alexa Fluor or HRP-conjugated secondary antibodies (**Table 4**) for 1 hour at RT. Images were acquired using the iBrightFL1000 system (Invitrogen). Signals were detected by fluorescence or chemiluminescence (Thermo Scientific, Cat# 34577).

### Identification of intracellular interaction partners by proximity biotinylation / nano liquid chromatography coupled to tandem mass spectrometry (nanoLC-MS/MS)

#### Sample Processing

HEK293T cells were transduced with lentivirus to stably overexpress wild-type (WT) GPR133-BioID2, mutant (H543R/T545A) GPR133-BioID2 or an empty vector control. Cells were treated with 50 μM biotin (Sigma, Cat# B4639-500MG) for 16 hours and whole cell lysates were prepared as described above. Biotinylated proteins were isolated using Pierce™ NeutrAvidin™ Agarose beads (Thermo Fisher, Cat# 29200) following the manufacturer’s protocol. Affinity-enriched proteins were separated by SDS-PAGE until the dye front entered 3 cm into the separating gel, and resulting gels were washed 3 times in distilled deionized H_2_O for 15 minutes each and visualized by staining overnight with EZ-Run Protein Gel Staining Solution (Thermo Fisher Scientific, Cat# BP36201). Stained protein gel regions were typically excised into 4 gel sections per gel lane, and destained as described (Xu et al., 2021). In-gel digestion was performed overnight with mass spectrometry grade trypsin (Trypsin Gold, Promega, Cat# V5280) at 5 ng/μL in 50 mM NH_4_HCO_3_ digest buffer. After acidification with 10% formic acid (final concentration of 0.5-1% formic acid), resulting peptides were desalted using hand-packed, reversed phase Empore C18 Extraction Disks (3M, Cat#3M2215), following an established method (Rappsilber et al., 2007).

#### Mass spectrometry data acquisition

Desalted peptides were concentrated to a very small droplet by vacuum centrifugation and reconstituted in 10 μL 0.1% formic acid in H_2_O. Approximately 90% of the peptide material was used for liquid chromatography, followed by tandem mass spectrometry (LC-MS/MS). A Q Exactive HF mass spectrometer was coupled directly to an EASY-nLC 1000 (Thermo Fisher Scientific, Cat#LC120) equipped with a self-packed 75 μm x 20-cm reverse phase column (ReproSil-Pur C18, 3M, Dr. Maisch GmbH, Germany) for peptide separation. Analytical column temperature was maintained at 50 °C by a column oven (Sonation GmBH, Germany). Peptides were eluted with a 3-40% acetonitrile gradient over 110 min at a flow rate of 250 nL/min. The mass spectrometer was operated in DDA mode with survey scans acquired at a resolution of 120,000 (at m/z 200) over a scan range of 300-1750 m/z. Up to 15 of the most abundant precursors from the survey scan were selected with an isolation window of 1.6 Th for fragmentation by higher-energy collisional dissociation with normalized collision energy (NCE) of 27. The maximum injection time for the survey and MS/MS scans was 60 ms and the ion target value (Automatic Gain Control) for both scan modes was set to 3e^6^.

#### Mass Spectrometry Data Processing

The mass spectra files were processed using the MaxQuant proteomics data analysis workflow (version 1.6.0.1) with the Andromeda search engine (Cox et al., 2011; Tyanova et al., 2016). Raw mass spectrometry files were used to extract peak lists which were searched with the Andromeda search engine against the human proteome and a file containing contaminants, such as human keratins. Trypsin digestion was specified allowing up to 2 missed cleavages with the minimum required peptide length set to be seven amino acids. N-acetylation of protein N-termini, oxidation of methionines and deamidation of asparagines and glutamines were set as variable modifications. For the initial identification search, parent peptide masses were allowed mass deviation of up to 20 ppm. Peptide spectral matches and protein identifications were filtered using a target-decoy approach at a false discovery rate of 1%. We used the raw MS1 intensity for protein quantification.

### Computational analysis of biotinylated interaction partners

All protein intensity values were log_10_ transformed and average log_10_intensities of empty vector controls (n=3) were subtracted from the values of both wild-type (WT, n=3) and mutant (H543R/T545A, n=3) GPR133 groups. For differential analysis, the proteins were then ranked by p value of unpaired t test between WT and mutants. To identify interactors shared by both WT and mutant GPR133, we set a p value threshold of p>0.9 for the WT-mutant comparison. The p value threshold was set at p<0.1 for differentially detected interactors. The raw mass spectrometry data and accompanying tables generated in this study are available at MassIVE (UCSD, https://massive.ucsd.edu/ProteoSAFe/static/massive.jsp) under accession number MSV000091163.

### Affinity purification of Strep-tagged GPR133 and co-purification of ESYT1

For input samples, whole cell lysates were prepared as described above. Twin-Strep-tagged GPR133 was purified using Strep-Tactin® XT coated magnetic beads (MagStrep “type3” XT Beads, IBA, Cat# 2-4090-002), according to the manufacturer’s protocol. In brief, after adding Biolock (IBA, Cat# 2-0205-250), whole cell lysates with incubated with MagStrep “type3” XT Beads overnight at 4°C. The next day, beads were collected with a magnetic separator and washed 3 times with 1x Buffer W (IBA, Cat# 2-1003-100). Proteins were eluted with 1X biotin elution buffer BXT (IBA, Cat# 2-1042-025). Laemmli buffer with DTT was added and elution samples were analyzed by Western blot as described above. Membranes were stained with a rabbit antibody specifically recognizing the cytosolic aspect of GPR133 (Sigma, Cat# HPA042395) or a rabbit antibody against ESYT1 (Sigma, Cat# HPA016858), and a goat-anti GAPDH antibody (Thermo Firsher, Cat# PA1-9046) as loading control for each membrane.

### Immunofluorescent staining and Proximity Ligation Assays

For immunofluorescent staining, cells were cultured on dishes coated with poly-L-ornithine and laminin, as described above, fixed with 4% paraformaldehyde (PFA, Sigma, Cat# P6148) for 20 min at RT, and blocked with 10% BSA in phosphate-buffered saline (PBS) for 1 hour at RT. Cells were then incubated with a primary antibody (**Table 4**) in 1% BSA in PBS at 4 °C overnight. The next day, cells were washed with PBS and stained with a secondary antibody (**Table 4**) for 1 hour at RT. Nuclei were stained with 500 ng/mL 4′,6-diamidino-2-phenylindole (DAPI) or Hoechst 333442 for 10 min at RT. For permeabilized staining, 0.1% Triton X-100 was added to the BSA- or PBS solution. For proximity ligation assays (PLA), cells were fixed, blocked and treated with primary antibodies from two different species at 4°C overnight. To detect proteins in close proximity, cells were then treated with PLUS and MINUS probes (Duolink® In Situ PLA® Probe Anti-Rabbit PLUS, Sigma, Cat# DUO92002 and Duolink® In Situ PLA® Probe Anti-Mouse MINUS, Sigma, Cat#DUO92004) followed by ligation and amplification steps according to the manufacturer’s protocol (Duolink® In Situ Detection Reagents Red, Sigma, DUO92008). For experiments involving intracellular Ca^2+^ increases, cells were treated with 1 μM thapsigargin (TG) (Millipore Sigma CAS 67526-95-8) or DMSO in DMEM for 2 min or 30 min at 37°C prior to fixation. Microscopy was conducted on a Zeiss Axiovert epifluorescent microscope or a Zeiss LSM700 laser scanning confocal microscope. To quantify PLA signals, a total of 2 - 5 fields were acquired in each experimental condition and PLA-positive spots were counted in relation to DAPI-positive cells. Fields were averaged for each biological replicate.

### Non-Permeabilized Enzyme-Linked Immunosorbent Assay (ELISA)

Cells were seeded onto 96-well plates coated with poly-L-ornithine and laminin, as described above. Twenty-four hours after seeding, cells were washed once with cold HBSS +Ca^2+^/+Mg^2+^ (Thermo Fisher, Cat# 14025092), fixed with 4% PFA for 20 min at RT, washed three times with PBS (Gibco, Cat# 14190-250) and blocked with DMEM containing 10% FBS for one hour at RT. Cells were then incubated with 8E3E8, a monoclonal mouse antibody specifically binding the N-terminus of GPR133 (Frenster *et al*., 2020; Frenster *et al*., 2021; Stephan *et al*., 2022), in DMEM containing 10% FBS. Cells were washed three times with PBS and incubated with horseradish peroxidase (HRP)-conjugated secondary antibodies (1:1000, chicken-anti mouse IgG Invitrogen Cat# A15975) in DMEM containing 10% FBS for 1 hour at RT. After three additional washes with PBS, cells were incubated with TMB (3,3’, 5,5’-tetramethylbenzidine)-stabilized chromogen (Thermo Fisher, Cat# SB02) for 5-10 min. The reaction was stopped by adding an equal volume of acidic stop solution (Thermo Fisher, Cat# SS04) and optical density/absorbance was measured at 450 nm (A_450 nm_).

### Homogeneous Time Resolved Fluorescence (HTRF)-based cAMP assays

HEK293T or GBM cells were seeded onto 96-well plates, pretreated with poly-L-ornithine and laminin, at a density of 75,000 cells (HEK293T) or 100,000 cells (GBM) per well. Twenty-four hours after seeding, cell culture medium, supplemented with 1 mM 3-isobutyl-1-methylxanthine (IBMX, Sigma-Aldrich, Cat# I7018-100MG) was added and cells were incubated at 37 °C for 30 – 60 min. In experiments involving elevated intracellular Ca^2+^, 1μM TG or DMSO were added to DMEM in addition to IBMX, also for a 30 min incubation at 37°C. Concentrations of cAMP were measured using the cAMP Gs dynamic kit (CisBio, Cat# 62AM4PEC), according to the manufacturer’s protocol.

### Fluorescent Ca^2+^ imaging

HEK293T cells were transduced with a lentiviral vector expressing the genetically encoded Ca^2+^ indicator GCaMP6s and the fluorescent protein TdTomato (Berlin et al., 2015; Liberti et al., 2016). They were imaged for 20-30 minutes on an Olympus IX73 epifluorescent microscope equipped with a SCMOS camera. Imaging data were analyzed on Olympus software.

### Tumorsphere formation assays

Patient-derived GBM cultures were cultured as described above. Cultures were dissociated into single cells and plated at 500 cells per well in a 96-well plate. Each experimental condition was carried out in 10 technical replicate wells. Cells were grown and supplemented with fresh media and growth factors for two weeks. Individual wells of 96-well plates were imaged using automated tile scanning on an EVOS imaging system (Thermo Fisher Scientific). Tile scans of each well were exported, and sphere numbers were counted using ImageJ.

### Visualization of RNA-seq data

RNA-seq data from HEK293 cells were downloaded from a previous publication (Aktas *et al*., 2017). The data were visualized on IGV (Integrative Genomics Viewer; https://igv.org/).

### Statistical analysis

All experiments were performed in biological replicates of at least three repeats (n > 3). Statistical analysis was performed using GraphPad Prism (version 8.4.3). Statistics are represented as mean ± standard error of the mean (SEM) as indicated. Statistical significance was calculated using either Students t-test, logrank test (for Kaplan-Meier survival curves), one-way analysis of variance (ANOVA), or two-way ANOVA, with Tukey’s or Sidak’s *post hoc* test for multiple comparisons. P values <0.05 were considered statistically significant (*p<0.05; **p<0.01; ***p<0.001; ****p<0.0001).

## Graphical abstract

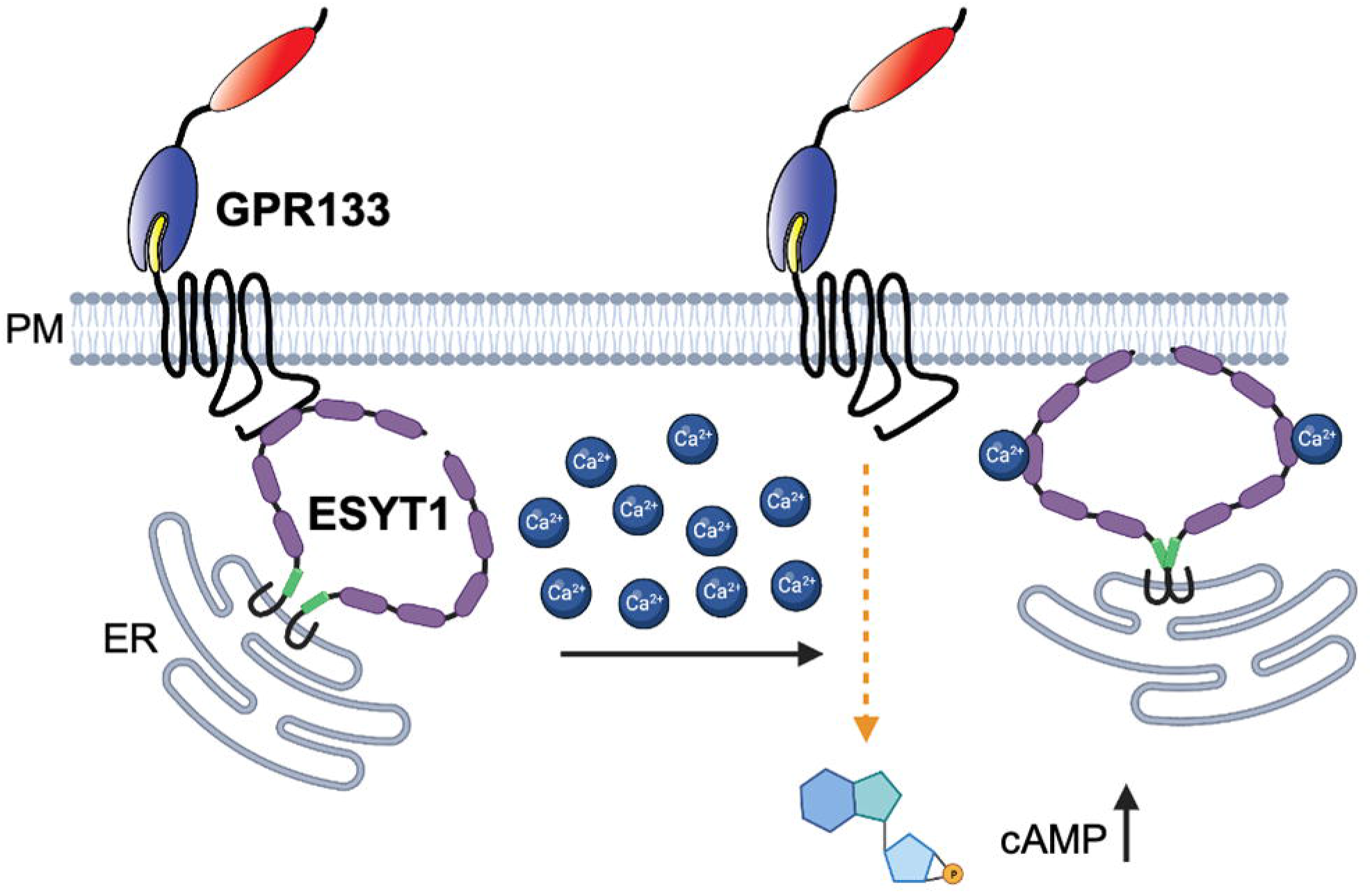

