## supplemental figures and legends for "Modulation of GPR133 (ADGRD1) Signaling by its Intracellular Interaction Partner Extended Synaptotagmin 1 (ESYT1)"

### Supplemental Information

**Supplementary Figure 1: Quality controls of HEK293T cells stably overexpressing GPR133-BioID2 and GPR133(H543R/T545A)-BioID2.**

(**A**) Immunofluorescent staining of HEK293T cells transduced with vector, GPR133-BioID2 or GPR133(H543R/T545A)-BioID2 using an anti-GPR133 antibody (red).

(**B**) Cellular cAMP levels were significantly increased in cells transduced with GPR133-BioID2, compared to the vector control or H543R/T545A-BioID2 (one-way ANOVA $F_{(2,6)}=15.29$, $p=0.0044$; Tukey's *post hoc* test: GPR133-BioID2 compared to vector, $p=0.0059$; GPR133-BioID2 compared to H543R/T545A, $p=0.0092$). Bars represent mean ± SEM of 3 experiments. ns, not significant.

(**C**) Overexpression of GPR133-BioID2 and GPR133 (H543R/T545A)-BioID2 is shown by Western blots of whole cell lysates. Staining with an antibody against GPR133's cytosolic C-terminus (C-term) detected bands corresponding to the cleaved and BioID2-tagged CTF (blue arrow, ~60 kDa) following transfection of GPR133-BioID2. Bands corresponding to the full-length receptor (red arrows, ~110 kDa) were detected following transfection of both GPR133-BioID2 (only a minor species, since the vast majority is cleaved) and the uncleavable H543R/T545A-BioID2 (essentially all protein is uncleaved). Staining with an antibody against biotin detected several bands in our whole cell lysates, including the bands corresponding to the CTF (blue arrow, ~60 kDa) and full-length receptor (red arrows, ~110 kDa), following transfection of GPR133-BioID2 and H543R/T545A-BioID, respectively.

(**D**) Samples were analyzed by Western blot and staining with an antibody against biotin before (input) and after (elution) neutravidin purification.

**Supplementary Figure 2: Biotinylated proteins with different intensities (empty vector subtracted) purified from HEK293T cells overexpressing wild-type GPR133 or GPR133 (H543R/T545A).**

Differentially biotinylated proteins between the HEK293T-GPR133-BioID2 and HEK293T-H543R/T545A-BioID2 conditions ($p<0.001$).

**Supplementary Figure 3: Confocal images of non-permeabilized HEK293T cells expressing GPR133 and ESYT1-GFP.**

Projections of Z stacks and orthogonal views are displayed. GPR133 (red) was detected with an antibody against its NTF. Intrinsic GFP fluorescence was used to identify ESYT1-GFP. Note that cell surface immunostaining of GPR133 is preserved whether the cells overexpress ESYT1-GFP or not. Nuclei were counterstained with Hoechst dye.

**Supplementary Figure 4: Bulk RNA-seq data from HEK293T cells.**

(**A**) Structural domains of ESYT1-3. All ESYTs share the first two C2 (C2A and C2B) domains. The C2E domain of ESYT1 is analogous to the C2C domains of ESYT2 and ESYT3, which mediate the tethering to PM phospholipids. The C2C and C2D domains of ESYT1 are unique and are thought to have arisen from duplication of C2A and C2B. The $Ca^{2+}$-binding capacity of C2C is considered necessary for the $Ca^{2+}$-dependent ER-PM tethering by ESYT1.

(**B**) RNA-seq tracks indicate that *ESYT1* and *ESYT2* are transcribed at similar levels in HEK293 cells, contrary to *ESYT3* (Aktas *et al*, 2017). The y axis (number of reads) is the same for all three tracks.

**Supplementary Figure 5: Western blot analysis confirms stable expression of GPR133 and overexpression of different ESYT1 constructs in whole cell lysates from HEK293T cells.**

Staining with anti-GPR133 (C-terminus) detects the same levels of GPR133 in all HEK293T cells stably expressing GPR133 (GPR133). No signal is detected in control HEK293T cells (CTRL). Staining with an anti-ESYT1 antibody detects endogenous ESYT1 in all samples. ESYT1 expression levels are increased following transfection of HEK293T cells (CTRL) and GPR133-expressing (GPR133) HEK293T cells with wild-type ESYT1, ∆C2C, ∆C2E or ∆C2C+E. Size shifts of ESYT1-specific bands correspond to the deletions of the C2C or C2E domain (ESYT1 ~ 123 kDa; ∆C2C ~109 kDa; ∆C2E ~ 109 kDa; ∆C2C+E ~ 95 kDa).

**Supplementary Figure 6: Effect of thapsigargin (TG) on intracellular $Ca^{2+}$.**

HEK293T cells expressing GCaMP6s were imaged for 30 min in control conditions (left) or after addition of 1 μM TG. Note the rapid increase in intracellular $Ca^{2+}$ ($[Ca^{2+}]_i$), as inferred from the increase in GCaMP6s fluorescence, after addition of TG, and its subsequent slow decay. Each trace represents the time course of a single cell in the depicted fields.

**Supplementary Figure 7: Effect of ESYT1 D724A on GPR133-expressing cells.**

(**A**) GPR133 surface expression following transfection of control HEK293T cells and HEK293T cells stably expressing GPR133 with wild-type ESYT1 or the point mutant ESYT1 D724A. Bars represent mean ± SEM of 4 to 8 experiments. Overexpression of ESYT1 or D724A did not affect GPR133 surface expression compared to the vector control (two-way ANOVA, $p>0.05$). $A_{450\ nm}$, absorbance/optical density at 450 nm.

(**B**) Intracellular cAMP levels following transfection of HEK293T cells stably expressing GPR133 with wild-type ESYT1 or the point mutant ESYT1 D724A. Bars represent mean ± SEM of 3 to 8 experiments. Concentrations of cAMP were significantly decreased in GPR133 expressing cells after transfection with ESYT1 and D724A compared to the vector control (two-way ANOVA $F_{(2,26)}=15.00$, $p<0.0001$; Sidak's *post hoc* test: GPR133 + vector vs. GPR133 + ESYT1, $p<0.0001$; GPR133 + vector vs. GPR133 + D724A, $p<0.0001$).
